# Calcium regulation of the *Arabidopsis* Na^+^/K^+^ transporter HKT1;1 improves seed germination under salt stress

**DOI:** 10.1101/2023.08.17.553671

**Authors:** Ancy E.J. Chandran, Aliza Finkler, Tom Aharon Hait, Yvonne Kiere, Sivan David, Metsada Pasmanik-Chor, Doron Shkolnik

## Abstract

Calcium is known to improve seed-germination rates under salt stress. We investigated the involvement of calcium ions (Ca^2+^) in regulating *HIGH-AFFINITY K^+^ TRANSPORTER 1* (*HKT1;1*), which encodes a Na^+^/K^+^ transporter, and its post-translational regulator *TYPE 2C PROTEIN PHOSPHATASE 49* (*PP2C49*), in germinating *Arabidopsis thaliana* seedlings. Germination rates of *hkt1* mutant seeds under salt stress remained unchanged by CaCl_2_ treatment in wild-type *Arabidopsis*, whereas *pp2c49* mutant seeds displayed improved salt-stress tolerance in the absence of CaCl_2_ supplementation. Analysis of *HKT1;1* and *PP2C49* promoter activity revealed that CaCl_2_ treatment results in radicle-focused expression of *HKT1;1* and reduction of the native radicle-exclusive expression of *PP2C49*. Ion-content analysis indicated that CaCl_2_ treatment improves K^+^ retention in germinating wild-type seedlings under salt stress, but not in *hkt1* seedlings. Transgenic seedlings designed to exclusively express *HKT1;1* in the radicle during germination displayed higher germination rates under salt stress than the wild type in the absence of CaCl_2_ treatment. Transcriptome analysis of germinating seedlings treated with CaCl_2_, NaCl, or both revealed 118 upregulated and 94 downregulated genes as responsive to the combined treatment. Bioinformatics analysis of the upstream sequences of CaCl_2_–NaCl-treatment-responsive upregulated genes revealed the abscisic acid response element CACGTGTC, a potential CaM-binding transcription activator-binding motif, as most prominent. Our findings suggest a key role for Ca^2+^ in mediating salt-stress responses during germination by regulating genes that function to maintain Na^+^ and K^+^ homeostasis, which is vital for seed germination under salt stress.

## INTRODUCTION

High concentrations of salt (NaCl) in the growth medium impair seed germination and seedling establishment. Increasing soil salinization of arable lands worldwide (Wang et al., 2003; Hassani et al., 2021) calls for a better understanding of the mechanisms that govern plant responses to salinity during early stages of development. Improvement of crop plant performance under salt-stress conditions is essential to ensuring food security and attenuating food scarcity (Tester and Langridge, 2010). Plants have evolved various strategies to adjust their growth and development to saline environments (Zhu, 2002), including removal of sodium ions (Na^+^) from the cytosol to the vacuole and the apoplast, mainly facilitated by the tonoplast-localized Na^+^/H^+^ antiporter NHX1 (Apse et al., 1999) and the plasma membrane-localized Na^+^/H^+^ antiporter SALT OVERLY SENSITIVE1 (SOS1) (Shi et al., 2000). Another type of salt response-related transporter in plants is the high-affinity K^+^-transporter (HKT) which facilitates long-distance Na^+^ transport to maintain Na^+^-level homeostasis at the whole-plant level (Waters et al., 2013). In *Arabidopsis*, a single HKT-type transporter (HKT1;1, AT4G10310) was identified and localized to the plasma membrane of the vascular parenchyma cells surrounding the xylem (Sunarpi et al., 2005; Davenport et al., 2007). In roots of young and mature *Arabidopsis*, *HKT1;1* functions in Na^+^ retrieval from the shootward xylem stream to adjacent parenchyma cells, thus limiting the accumulation of Na^+^ in salt-sensitive shoot tissues (Mäser et al., 2002; Rus et al., 2004; Sunarpi et al., 2005; Horie et al., 2006; Møller et al., 2009; Shkolnik-Inbar et al., 2013; Möller et al., 2017). Furthermore, *hkt1* mutants, which are characterized as salt-hypersensitive, accumulate higher levels of Na^+^ in the shoots and xylem vessels (Mäser et al., 2002; Sunarpi et al., 2005; Davenport et al., 2007). However, the mechanisms by which these and other transport systems function to maintain Na^+^ homeostasis during germination in association with other salt stress-related signaling pathways and responses have yet to be fully investigated.

Calcium ion (Ca^2+^) has been shown to promote salt tolerance of *Arabidopsis*, common reed and cotton (Kent and Läuchli, 1985; Zehra et al., 2012; Shkolnik et al., 2019). Ca^2+^ has been proposed to improve the selectivity of potassium ion (K^+^) uptake to maintain K^+^/Na^+^ ratios upon exposure to excessive concentrations of NaCl (LaHaye and Epstein, 1969; Bernstein, 1970; Lahaye and Epstein, 1971). Ca^2+^ is a ubiquitous secondary messenger that mediates a wide array of downstream signaling cascades in response to various internal and environmental stimuli, including salt (Choi et al., 2014; Choi et al., 2016; Shkolnik et al., 2018). In roots, NaCl induces elevation of free Ca^2+^ levels in the cytosol by promoting Ca^2+^ influx from the vacuole, as in the activity of the tonoplast-localized TWO PORE CHANNEL1 (TPC1) (Peiter et al., 2005; Evans et al., 2016). Moreover, other subcellular compartments that function as Ca^2+^ reservoirs, such as the endoplasmic reticulum, chloroplasts, mitochondria and peroxisomes, may also be involved (Stael et al., 2012; Choi et al., 2014). In plant cells, Ca^2+^ signals are perceived by multiple calmodulin (CaM) and CaM-like proteins, and the duration, localization, intensity and frequency of the cytosolic Ca^2+^ transients determine the downstream responses, involving the activity of various cellular components, such as phosphatases, kinases and transcription factors (TFs) (Kudla et al., 2010). The SOS signaling pathway is particularly activated in response to NaCl. The myristoylated Ca^2+^-binding protein SOS3 functions as a primary Ca^2+^ sensor that perceives the signals and interacts with the serine/threonine protein kinase SOS2, which in turn activates SOS1 to facilitate Na^+^ removal from the cytosol to the apoplast (Qiu et al., 2002; Ji et al., 2013). Nevertheless, the molecular mechanism underlying the Ca^2+^-mediated improved tolerance to salt stress in general, and during germination in particular, has yet to be deciphered.

*Arabidopsis* is a powerful tool for investigating molecular mechanisms of plant responses to salt stress (Zhu, 2000; Yang and Guo, 2018). Investigation of the involvement of Ca^2+^ signaling in response to abiotic stress in *Arabidopsis* revealed six genes encoding CaM-binding transcription activators (CAMTAs), all with similar DNA-binding domains (Bouché et al., 2002; Yang and Poovaiah, 2002; Mitsuda et al., 2003; Finkler et al., 2007a; Doherty et al., 2009; Shen et al., 2015). CAMTA proteins were suggested to be components of Ca^2+^-mediated signaling pathways in plant responses to various external stimuli, including abiotic stresses and hormones such as abscisic acid (ABA) (Yang and Poovaiah, 2002; Finkler et al., 2007a). CAMTA-similar genes were then identified in other plant species, including tomato (*Solanum lycopersicum*; Li et al., 2014), maize (*Zea mays*; Yue et al., 2015), rice (*Oryza sativa*; Choi et al., 2005; Rahman et al., 2016), and more. Recently, CAMTA6 has been shown to regulate the seed’s response to salt stress during germination by affecting the expression of multiple genes, and examination of presumed promoters of salt-responsive CAMTA6-dependent upregulated genes revealed the high frequency of the ABA response element (ABRE) CACGTGTC, suggested as a potential CAMTA-binding site (Shkolnik et al., 2019). In the background of a *camta6* mutation, *HKT1;1* was expressed predominantly in the radicle (Shkolnik et al., 2019). This unique radicle-focused expression pattern of *HKT1;1* was associated with the *camta6* phenotype of improved germination in the presence of salt, which was not observed in *camta6* seedlings that were several days old and found to display salt hypersensitivity (Shkolnik et al., 2019). This could be explained by the fact that in pre-germinating seedlings, in which the vascular tissues have not yet differentiated, *HKT1;1* was evenly expressed in all tissues (Shkolnik et al., 2019). Hence, the activity and influence of HKT1;1 on systemic Na^+^-level regulation in germinating seedlings differ from its function of limiting shoot Na^+^ accumulation in later stages of development (Davenport et al., 2007). Recently, HKT1;1 has been reported to function in protecting plant fertility by limiting Na^+^ accumulation in *Arabidopsis* stamen tissues (Uchiyama et al., 2023) and in tomato, HKT1;2 was shown to regulate Na^+^/K^+^ homeostasis in salt-stressed shoots (Jaime-Pérez et al., 2017). Overexpression of the CaM-like protein CML40 in *Medicago truncatula* led to downregulation of *MtHKT1;1* and *MtHKT1;2* expression, which resulted in hypersensitivity to salt during germination (Zhang et al., 2019). In wheat (*Triticum aestivum*), *HKT1* silencing by antisense RNA resulted in improved tolerance to NaCl, which was attributed to reduced Na^+^ uptake (Laurie et al., 2002), in agreement with the proposed function of both HKT1 from wheat and HKT1;1 from *Arabidopsis* in facilitating Na^+^ entry into plant cells (Gassmann et al., 1996; Uozumi et al., 2000). Furthermore, *HKT1;1* was found to be negatively regulated by the ABA- and salt-regulated TF *ABA INSENSITIVE 4* (Finkelstein, 1994; Shkolnik and Bar-Zvi, 2008; Shkolnik-Inbar et al., 2013), which is consistent with the salt hypersensitivity of *hkt1* seeds (Shkolnik et al., 2019). Recently, *HKT1;1* was found to be directly inactivated by the phosphatase PP2C49 (At3G62260) (Chu et al., 2021), a member of the type 2C protein phosphatase family which facilitates dephosphorylation of substrate proteins that are involved in signaling in *Arabidopsis* (Luan, 2003; Schweighofer et al., 2004). Chu et al. (2021) depicted *PP2C49* expression in almost all tissues of juvenile and mature *Arabidopsis*, excluding germinating seedlings, and specifically in the root vasculature, overlapping with the expression pattern of *HKT1;1*. Nevertheless, the regulation of HKT1;1 by PP2C49 during germination, under control and salt stress, has not been studied.

Here, we found that Ca^2+^ promotes downregulation of *PP2C49* to enable HKT1;1-induced improvement of K^+^ retention, and we provide further insights into the effect of Ca^2+^ on the transcriptome profile of germinating seedlings under salt stress. Moreover, transcriptome analysis indicated that genes functioning in ABA signaling and members of the CAMTA and teosinte branched1/cincinnata/proliferating cell factor (TCP) TF families are significant players in the Ca^2+^-mediated response to salt stress during germination.

## RESULTS

### Germinating *hkt1* mutants are not responsive to CaCl_2_ treatment under salt stress

Seeds of *hkt1* mutants are hypersensitive to salt during germination (Shkolnik et al., 2019). We examined the germination rate of seeds of two allelic *hkt1* mutants (*hkt1* and *hkt1-4*) and the wild type (*Arabidopsis* Col-0) sown on agar-solidified medium supplemented with 0 (control), 150 mM or 200 mM NaCl. Germination was considered only when green, fully unfurled green cotyledons were observed. When sown on 150 mM or 200 mM NaCl-containing medium, wild-type seeds displayed germination rates of 78.3% ± 3.0% and 2.6% ± 0.5%, respectively, in contrast to, respectively, 33.7% ± 7.3% and 1.3% ± 0.5% germination for *hkt1* and 35.3% ± 9.0% and 1.6% ± 0.5% germination of *hkt1-4* (Fig. 1). Conversely, when sown on mannitol-containing medium, no significant differences in the germination rates of wild type and *hkt1* mutants were scored (Supplemental Fig. S1), indicating that the *hkt1* salt hypersensitivity is not due to a general response to osmotic stress. Seeds of all genotypes displayed germination rates of ∼100% when no NaCl was added to the growth medium (Fig. 1). Germination medium supplemented with CaCl_2_ at 10 mM (1 mM in the control) in the presence of 150 mM or 200 mM NaCl improved germination rates, as previously demonstrated (Shkolnik et al., 2019) and confirmed here with the wild-type seeds, which displayed 58% ± 4% and 27% ± 7% germination, respectively (Fig. 1). Interestingly, germination of *hkt1* and *hkt1-4* seeds sown on 150 mM or 200 mM NaCl medium supplemented with CaCl_2_ was not improved—30.3% ± 9.7%, 4% ± 2% germination for *hkt1* and 31% ± 10%, 3.4% ± 1.5% germination for *hkt1-4*, respectively (Fig. 1).

**Figure 1.**
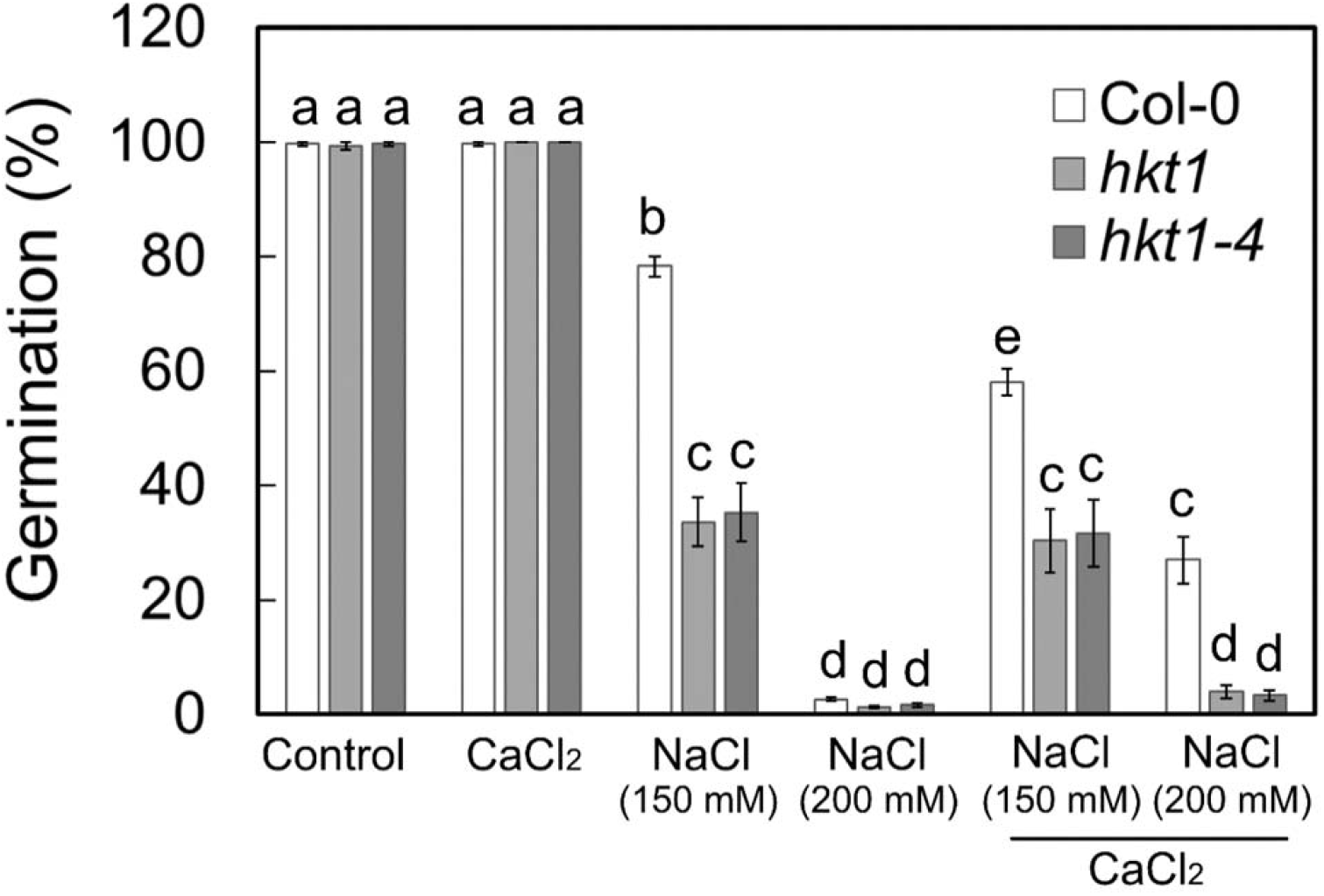
Germination of *hkt1* seeds under salt stress is unaffected by CaCl_2_ treatment. Seeds of the indicated genotypes were sown on agar-solidified 0.25X MS medium supplemented with the indicated single or combined chemicals (NaCl, 150 mM or 200 mM; CaCl_2_, 10 mM) and germination rates were scored 5 days after plating. Data are means ± SE (three independent biological experiments, ∼100 seeds each), and different letters above the bars indicate significantly different values by Tukey’s HSD post hoc test (*P* < 0.01).

To broaden our understanding of HKT1;1 function in the Ca^2+^-mediated response to salt stress, we examined the growth and development of roots of *hkt1* mutants at later stages of development (3-to 11-day-old seedlings, grown on agar-solidified medium; see Materials and Methods), supplemented with 0 (control), or 10 mM CaCl_2_, 200 mM NaCl, or both (Supplemental Fig. S2). Interestingly, supplementation of CaCl_2_ improved growth of the primary root of seedlings of all genotypes under salt-stress conditions; however, lateral-root count revealed reduced responsiveness of *hkt1* to the elevated CaCl_2_ concentration in the presence of NaCl, reflected by significantly reduced lateral-root formation (Supplemental Fig. S2). This indicated the involvement of HKT1;1 in the Ca^2+^-mediated response to salt at this developmental stage as well. No significant phenotypic differences were observed among the three genotypes under control conditions or in response to CaCl_2_ treatments (Supplemental Fig. S2).

### Ca^2+^ promotes radicle-focused expression of *HKT1;1*

To study the effect of CaCl_2_ treatment on the expression pattern of *HKT1;1*, we used transgenic seedlings expressing the GUS reporter gene under regulation of the *HKT1;1* promoter (*HKT1;1pro*:GUS) (Shkolnik et al., 2019). CaCl_2_ treatment resulted in radicle-focused *HKT1;1* promoter activity, which was enhanced in response to the combined CaCl_2_ and NaCl treatment (Fig. 2A), further indicating involvement of HKT1;1 in the Ca^2+^-mediated improvement of germination in the presence of salt. Under control conditions, *HKT1;1* was expressed evenly in all tissues of the germinating seedlings, and NaCl treatment resulted in enhanced expression in the whole seedling compared to controls (Fig. 2A) (Shkolnik et al., 2019). To further validate the observed *HKT1;1*-expression pattern in response to treatment with NaCl, CaCl_2_, or both, we performed RT-qPCR analysis for relative *HKT1;1* transcript level quantification, based on RNA that was isolated from whole seedlings, or separated cotyledon or radicle tissues (see Materials and Methods). In agreement with the promoter-activity analysis, transcripts of *HKT1;1* accumulated in response to CaCl_2_ treatment in the radicle (2.17 ± 0.33, relative to expression in the whole seedling which was set to a value of 1.0) and less so in the cotyledon (0.22 ± 0.08)—more pronounced in response to the combined CaCl_2_–NaCl treatment (radicle, 5.46 ± 0.65; cotyledon, 0.19 ± 0.05) (Fig. 2B). In fact, CaCl_2_ treatment, with or without NaCl supplementation, appeared to significantly reduce *HKT1;1* expression in the cotyledons (Fig. 2B). The previously reported enhanced whole-seedling expression in response to NaCl treatment (Shkolnik et al., 2019) was reproduced (radicle, 3.8 ± 0.65; cotyledon, 3.1 ± 0.55) (Fig. 2, A and B).

**Figure 2.**
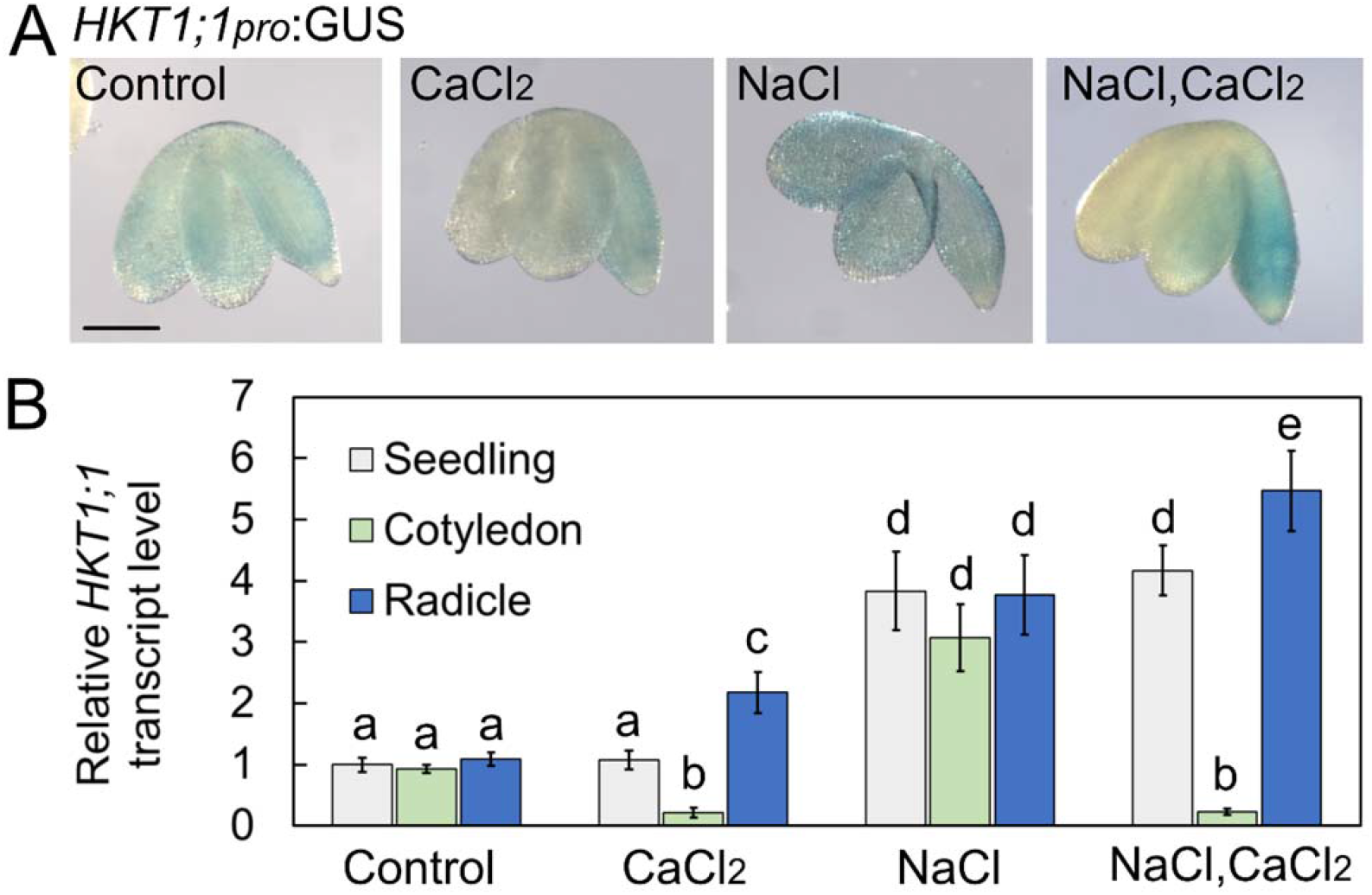
CaCl_2_ treatment of pre-germinating seedlings results in radicle-focused expression of *HKT1;1*. A, GUS staining of *HKT1;1pro*:GUS-harboring pre-germinating seedlings treated with single or combined chemicals as indicated (NaCl, 200 mM; CaCl_2_, 10 mM) for 16 h prior to ejection, staining and imaging. Bar = 1 mm. B, RT-qPCR analysis for relative *HKT1;1* transcript level quantification in pre-germinated seedlings. Data are means ± SD (three biological experiments, ∼30 seeds each) and different letters above the bars indicate significantly different values by Tukey’s HSD post hoc test (*P* < 0.01).

### Radicle-focused *HKT1;1* expression promotes salt tolerance during germination

To determine whether the Ca^2+^-mediated improvement in salt tolerance during germination is due to the observed diminution of *HKT1;1* expression in the cotyledons or to the radicle-focused expression, we created transgenic seedlings that express the coding sequence of *HKT1;1* under regulation of the native promoter of *Arabidopsis* AT4G31830. This gene has previously been reported to drive radicle-specific expression during germination (Jeong et al., 2014). The construct was introduced into the *hkt1-4* mutant to avoid native HKT1;1 activity and referred to as “HKT1;1 Radicle-expressed” (HKT1;1-RE). RT-qPCR analysis for *HKT1;1* relative transcript level quantification confirmed *HKT1;1* expression in the radicles in HKT1;1-RE germinating seedlings (radicle, 4.4 ± 1.3; cotyledon, 0.23 ± 0.1, relative to whole-seedling expression) (Fig. 3A). Equal relative transcript levels of *HKT1;1* were measured in wild-type tissues and very low expression levels were noted in all tissues of the *hkt1-4* mutant (Fig. 3A). These findings confirmed that the mutant can serve as a suitable background for tissue-specific expression of *HKT1;1*. Next, we examined germination of HKT1;1-RE seeds in the presence of 200 mM NaCl in the germination medium. Remarkably, seeds of HKT1;1-RE displayed 31 ± 5.6% germination under salt stress (Fig. 3B), similar to the germination rate of the wild-type in the presence of NaCl with CaCl_2_ supplementation (Fig. 1). Collectively, these findings strongly associate the radicle-enhanced expression of *HKT1;1*, rather than the diminished *HKT1;1* expression in cotyledons, with salt tolerance during germination.

**Figure 3.**
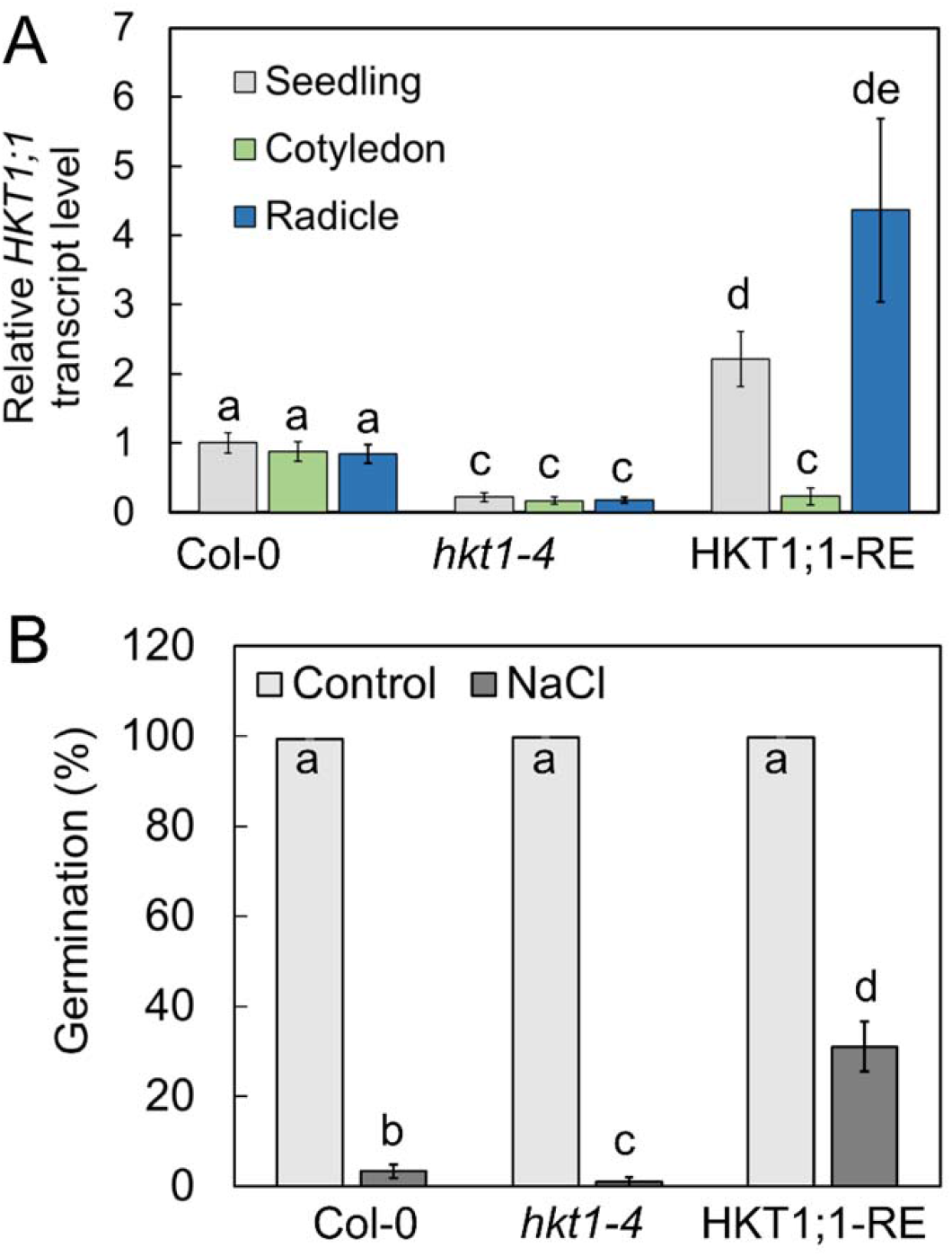
HKT1;1-RE seeds display improved germination under salt stress conditions. A, RT-qPCR analysis for *HKT1;1* relative transcript level quantification in wild-type (Col-0), *hkt1-4* mutant and HKT1;1-RE pre-germinated seedlings. Data are means ± SD (three biological experiments, ∼30 seeds each). B, Germination assay. Seeds of wild-type, *hkt1-4* mutant and HKT1;1-RE were sown on agar-solidified 0.25X MS medium supplemented with 0 (control) or 200 mM NaCl. Data are means ± SE (three independent biological experiments, ∼100 seeds each). Different letters above the bars indicate significantly different values by Tukey’s HSD post hoc test (*P* < 0.01).

### *hkt1* mutants accumulate less Na^+^ and more K^+^ during germination under salt stress

Plant cells function to limit the accumulation of toxic levels of Na^+^ in the cytosol and to promote retention of sufficient levels of K^+^ (Munns and Tester, 2008; Sun et al., 2015; Almeida et al., 2017). To quantify the accumulation of Na^+^ and K^+^ in wild-type and *hkt1* mutant pre-germinating seedlings, extracts of seedlings ejected from seeds that had been treated with 0 (control), or 10 mM CaCl_2_, 150 mM NaCl, or both for 16 h were analyzed for ion content by inductively coupled plasma optical emission spectrometry (ICP-OES) as previously described (Shkolnik et al., 2019). Under control conditions, wild-type and *hkt1* seedlings accumulated 1.39 ± 0.15 µg and 1.47 ± 0.1 µg Na^+^ per seedling, respectively, and in response to CaCl_2_ treatment, they accumulated 1.29 ± 0.14 µg and 1.37 ± 0.18 µg Na^+^ per seedling, respectively (Fig. 4). In the presence of NaCl, wild-type and *hkt1* seedlings accumulated 2.34 ± 0.28 µg and 2.87 ± 0.17 µg Na^+^ per seedling, respectively (Fig. 4), suggesting that the *hkt1* hypersensitivity to salt involves enhanced accumulation of Na^+^. In response to the combined NaCl–CaCl_2_ treatment, wild-type and *hkt1* seedlings accumulated 1.99 ± 0.12 µg and 3.1 ± 0.11 µg Na^+^ per seedling, respectively (Fig. 4), strongly indicating the involvement of HKT1;1 in maintaining Ca^2+^-regulated Na^+^ homeostasis in germinating seedlings under salt stress. Furthermore, NaCl treatment was found to significantly deplete the K^+^ content in both wild-type, as previously reported (Sun et al., 2015), and *hkt1* pre-germinating seedlings, with 0.56 ± 0.07 µg and 0.39 ± 0.15 µg K^+^ per seedling, respectively (Fig. 4), strongly suggesting the involvement of HKT1;1 in K^+^ retention under salt stress. Interestingly, in response to the combined NaCl–CaCl_2_ treatment, improved accumulation of K^+^ in wild-type seedlings was measured (1.04 ± 0.17 µg per seedling) but not in *hkt1* seedlings (0.42 ± 0.12 µg per seedling) (Fig. 4). Under control conditions, wild-type and *hkt1* seedlings accumulated an equal basal level of K^+^ of 2.6 ± 0.07 µg and 2.56 ± 0.15 µg per seedling, respectively (Fig. 4), and following CaCl_2_ treatment, they accumulated 2.61 ± 0.1 µg and 2.54 ± 0.24 µg per seedling, respectively (Fig. 4). Collectively, these data explain, at least in part, the hypersensitivity of *hkt1* seedlings to salt stress and their lack of responsiveness to the germination-improving effect of CaCl_2_ in the presence of salt.

**Figure 4.**
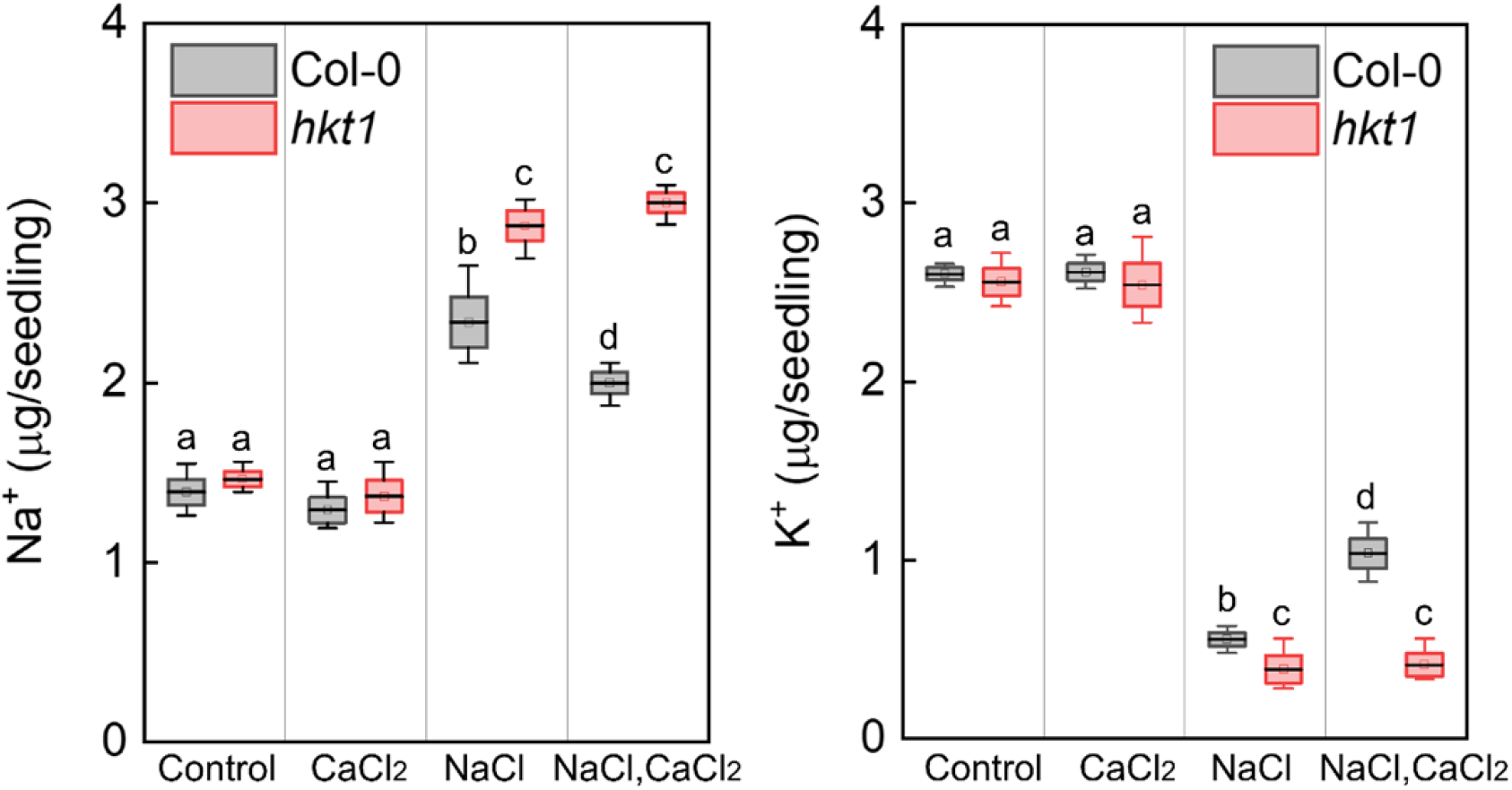
*hkt1* mutant accumulates more Na^+^ and less K^+^ under salt-stress conditions. Ion content was determined as described in Materials and Methods in pre-germinated wild-type (Col-0) and *hkt1*-mutant seedlings treated with 0 (control), 10 mM CaCl_2_, 150 mM NaCl or both, as indicated, for 16 h prior to seedling ejection. Data are mean values (black bar in box) ± SD (three biological experiments, ∼40 seeds each) and different letters above the bars indicate significantly different values by Tukey’s HSD post hoc test (*P* < 0.01).

### *pp2c49* mutants display tolerance to salt stress during germination

The phosphatase PP2C49 directly inhibits the activity of HKT1;1, and is associated with salt hypersensitivity in young *Arabidopsis* seedlings (Chu et al., 2021). To study the effect of Ca^2+^ supplementation on the germination rate of *pp2c49* mutant seeds under salt stress, we performed a germination assay with seeds of three *pp2c49* allelic mutants and the wild-type. Strikingly, when sown on 200 mM NaCl-containing medium, all three *pp2c49* allelic mutants displayed improved tolerance to NaCl, with germination rates of 28 ± 8.16%, 26.75 ± 5.7% and 41.5 ± 8.1% (*pp2c49-1, pp2c49-2* and *pp2c49-3*, respectively), compared to wild-type seeds with a germination rate of 2.6 ± 0.5% (Fig. 5A). Supplementation of CaCl_2_ to the NaCl-containing medium did not significantly affect the germination rate of *pp2c49* mutants, whereas it did for wild-type seeds (Fig. 5A). These results strongly suggest that PP2C49 is an important component in the Ca^2+^-regulated signaling pathway in response to salt stress during germination.

**Figure 5.**
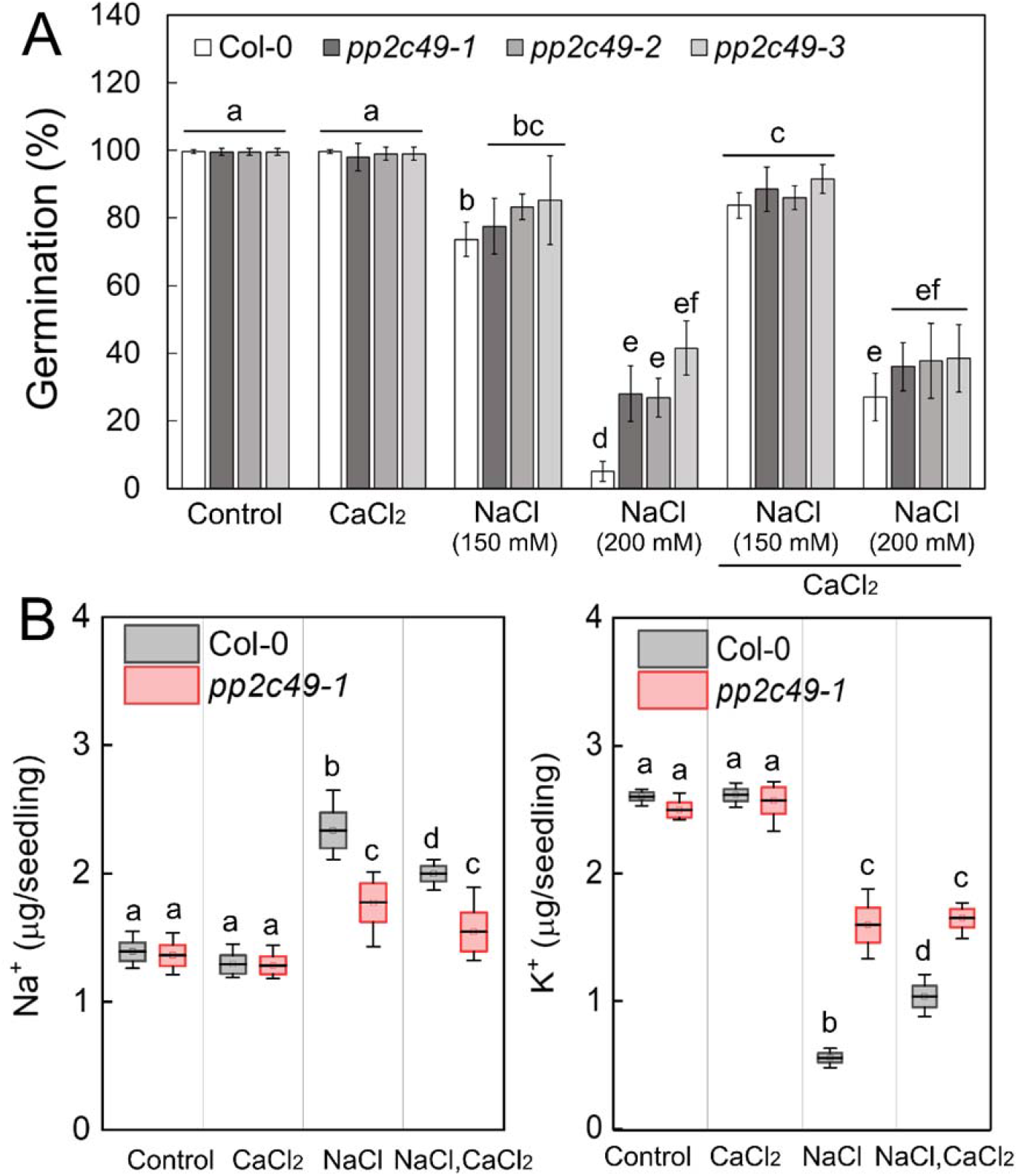
*pp2c49* mutants display improved tolerance to salt during germination as compared to wild type. A, Seeds of the indicated genotypes were sown on agar-solidified 0.25X MS medium supplemented with single or combined chemicals as indicated (NaCl, 150 mM or 200 mM; CaCl_2_, 10 mM), and germination rates were scored 5 days after plating. Data are means ± SE (three independent biological experiments, ∼100 seeds each). B, Ion content was determined as described in Materials and Methods in pre-germinated wild type (Col-0) and *pp2c49-1* mutant seedlings treated with 0 (control), 10 mM CaCl_2_, 150 mM NaCl or both, as indicated, for 16 h prior to seedling ejection. Data are mean values (black bar in box) ± SD (three biological experiments, ∼40 seeds each). Different letters above the bars indicate significantly different values by Tukey’s HSD post hoc test (*P* < 0.01).

To examine whether the *pp2c49* phenotype of improved tolerance to salt during germination is related to Ca^2+^ regulation of the accumulation of Na^+^, K^+^, or both in germinating seedlings, we quantified the ion content in a *pp2c49-1* mutant and the wild-type in response to treatment with NaCl, CaCl_2_, or both. In response to NaCl treatment, *pp2c49-1* pre-germinating seedlings accumulated 1.77 ± 0.3 µg Na^+^ per seedling, significantly less than the wild-type with 2.34 ± 0.28 µg Na^+^ per seedling (Fig. 5B). Furthermore, combining CaCl_2_ with NaCl did not reduce the Na^+^ accumulation of *pp2c49-1* (1.54 ± 0.3 µg per seedling) but it did reduce it in the wild type (1.99 ± 0.12 µg per seedling) (Fig. 5B). In response to NaCl treatment, wild-type and *pp2c49-1* pre-germinating seedlings accumulated 0.55 ± 0.07 µg and 1.6 ± 0.27 µg K^+^ per seedling, respectively, and in response to the combined NaCl–CaCl_2_ treatment, they accumulated 1.04 ± 0.17 µg and 1.65 ± 0.15 µg K^+^ per seedling, respectively; they accumulated similar levels of K^+^ in response to the CaCl_2_ treatment (Fig. 5B). Under control conditions and in response to the CaCl_2_ treatment, seedlings of both genotypes accumulated similar levels of Na^+^ as well (Fig. 5B). Collectively, these data strongly indicate the involvement of PP2C49 in Ca^2+^-mediated maintenance of ion homeostasis under salt stress during germination, which is consistent with the *pp2c49* phenotype of improved tolerance.

PP2C49 has been found to be expressed in almost all tissues of young and mature *Arabidopsis* (Chu et al., 2021). Nevertheless, its expression pattern in germinating seedlings has never been assessed. *PP2C49* promoter activity *in planta,* using *PP2C49pro*:GUS-expressing seedlings, in response to control, NaCl, CaCl_2_ or NaCl–CaCl_2_ treatments, indicated that *PP2C49* is exclusively expressed in the radicle under all conditions, and that this expression is most pronounced in response to the NaCl treatment (Fig. 6A). Moreover, CaCl_2_ treatment significantly reduced the radicle-confined *PP2C49* expression and abolished the inductive effect of NaCl on its expression (Fig. 6A). To reconfirm these data, we performed RT-qPCR for relative quantification of *PP2C49* transcript level in whole germinating seedlings, and cotyledon and radicle tissues (Fig. 6B). Indeed, CaCl_2_ significantly reduced the relative *PP2C49* transcript level in the whole seedling (0.43 ± 0.11), cotyledons (0.35 ± 0.08) and radicle (0.37 ± 0.21) compared to the control treatment, and abolished transcript accumulation in response to NaCl treatment when the seeds were exposed to the combined treatment—in the whole seedling (0.4 ± 0.19), cotyledons (0.26 ± 0.07) and radicle (0.33 ± 0.14)—in agreement with the expression pattern detected by GUS staining (Fig. 6). These data strongly indicate negative regulation of the expression of *PP2C49* by Ca^2+^ and consequently, attenuation of its negative regulation of HKT1;1, thereby leading to improved germination under salt stress.

**Figure 6.**
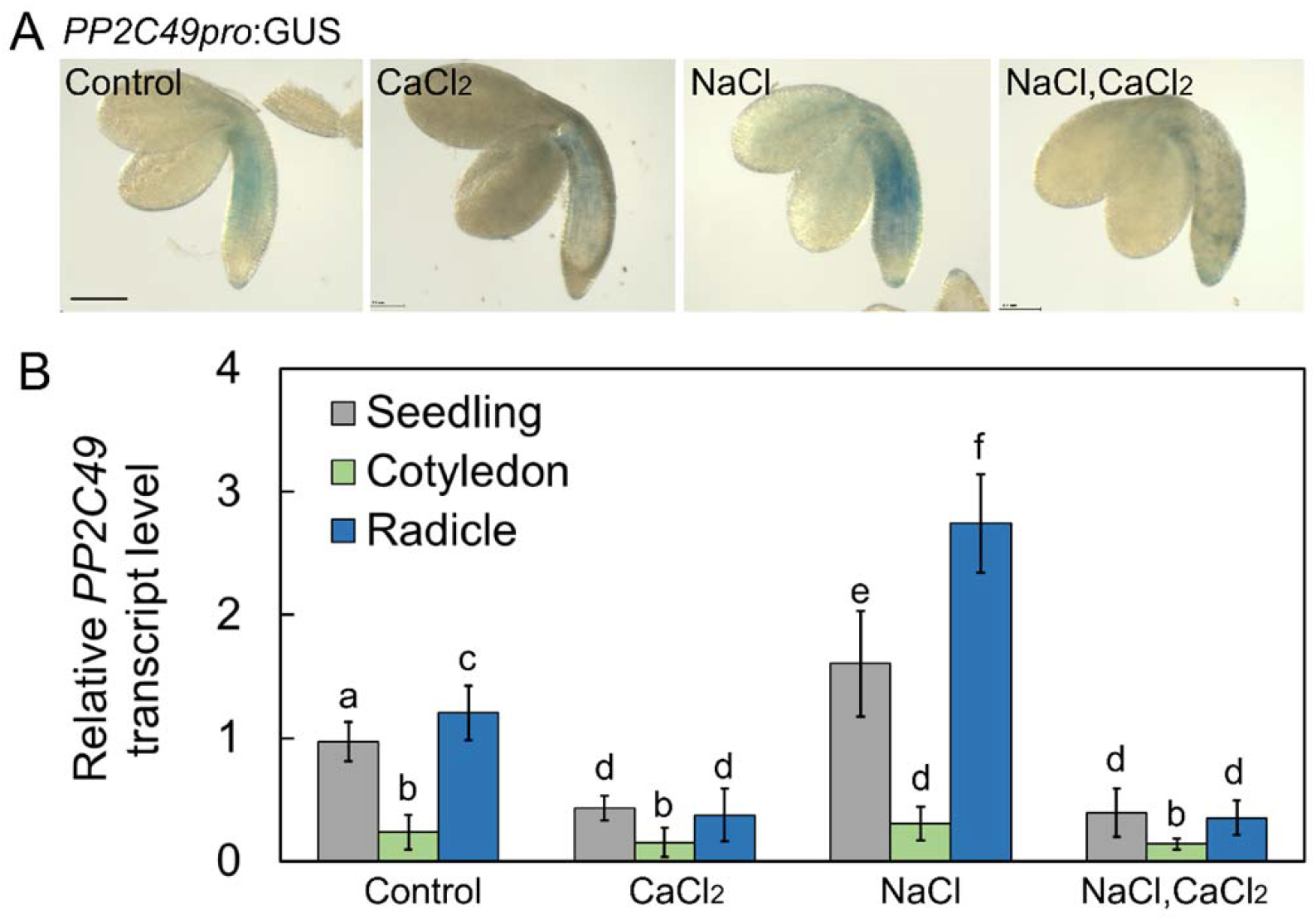
*PP2C49* is expressed in the radicles and downregulated by CaCl_2_ treatment. A, GUS staining of *PP2C49pro*:GUS-harboring pre-germinating seedlings treated with single or combined chemicals as indicated (NaCl, 200 mM; CaCl_2_, 10 mM) for 16 h prior to ejection, staining and imaging. Bar = 1 mm. B, RT-qPCR analysis for relative *PP2C49* transcript level quantification in pre-germinated seedlings. Data are means ± SD (three biological experiments, ∼30 seeds each) and different letters above the bars indicate significantly different values by Tukey’s HSD post hoc test (*P* < 0.01).

### Transcriptome analysis of CaCl_2_-treated germinating seedlings under salt stress

Transcriptome analysis was performed to further assess the extent of the effect of CaCl_2_ supplementation on germination under salt stress. RNA was isolated from pre-germinating seedlings that had been treated with CaCl_2_, NaCl, or both (see Materials and Methods). Expression reads higher than 10 were filtered for differential expression using Partek Genomics Suite v7.20.0831. Differentially expressed gene (DEG) lists of CaCl_2_-treated, NaCl-treated, and NaCl–CaCl_2_-treated seedlings versus the control treatment (cutoff fold change [FC] ≥ |1.25| and *P* ≤ 0.05) were extracted (Supplemental Tables S1–S3). Venn diagrams were created to compare DEG lists (cutoff FC ≥ |1.25| and *P* ≤ 0.05; Supplemental Fig. S3 and Table S4). Using these selected cutoffs, 118 unique upregulated and 94 unique downregulated NaCl–CaCl_2_-responsive genes were identified (Supplemental Table S4). In response to CaCl_2_ treatment, 856 unique upregulated and 195 unique downregulated genes were identified, whereas in response to NaCl treatment, 244 unique upregulated and 142 unique downregulated genes were identified (Supplemental Fig. S3 and Table S4). Functional analysis of these upregulated and downregulated genes, exclusive to each treatment (CaCl_2_, NaCl, CaCl_2_–NaCl) versus the control, was conducted using the DAVID Gene Functional Classification tool (https://david.ncifcrf.gov/) (Sherman et al., 2022), with multiple sources of functional annotations (https://david.ncifcrf.gov/content.jsp?file=update.html). Classifications were selected based on *P* values (*P* ≤ 0.05). Highly significantly enriched GO terms for differentially expressed (CaCl_2_–NaCl treatment versus control) upregulated and downregulated genes were of nuclear proteins (∼22% upregulated and ∼36% downregulated) and expression-related genes (∼44% upregulated), RNA binding (20% downregulated) and TFs (∼8.8% upregulated and ∼17% downregulated) (Fig. 7, Supplemental Table S5). Interestingly, ∼38% of the upregulated DEGs were classified as involved in the response to abiotic stress, similar to the response to NaCl treatment (Fig. 7, Supplemental Table S5). Furthermore, in response to NaCl treatment, most of the downregulated DEGs were of organellar proteins (∼82%), with ∼32% associated with the chloroplast and ∼11% to the nucleus, whereas ∼9% of the upregulated DEGs were associated with the nuclear lumen (Fig. 7, Supplemental Table S5). Similar to the response to NaCl treatment, ∼79% of the downregulated DEGs in response to CaCl_2_ treatment were organellar; however, ∼21% of the downregulated DEGs were associated with the nucleus, whereas ∼31% were of chloroplast proteins (Fig. 7, Supplemental Table S5). There was therefore a significant negative effect of NaCl and positive effect of CaCl_2_ on chloroplast-associated protein expression in germinating seedlings, which was not observed in response to the combined treatment. NaCl induced ABA-responsive DEGs (∼14%), as expected in view of the involvement of ABA in salt-responsive signaling pathways (Zhu, 2002). On the other hand, CaCl_2_ downregulated DEGs of this functional classification (∼12%) (Fig. 7, Supplemental Table S5). The full raw and selected DEG lists are available in Supplemental Table S5.

**Figure 7.**
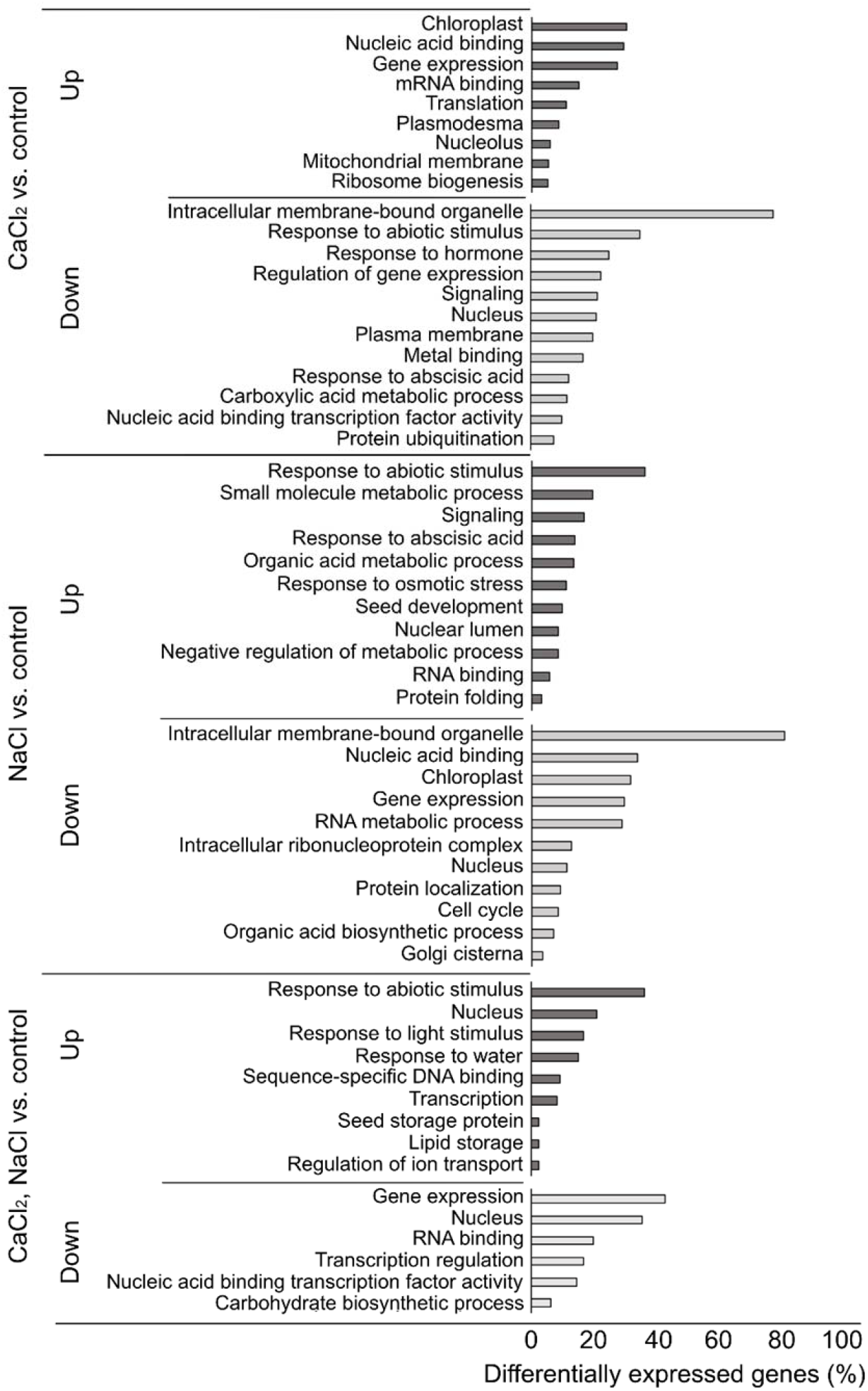
Functional classification of differentially expressed genes by the DAVID gene functional classification tool (https://david.ncifcrf.gov/). The tool utilizes the comprehensive DAVID Knowledgebase which pulls together multiple sources of functional annotations. Representative classifications were selected base on *P* values (*P* ≤ 0.05). Classification analysis of the upregulated and downregulated differentially expressed genes was performed based on the following: biological process, molecular function and cellular component.

### Promoter analysis of salt- and calcium-responsive genes: an unbiased motif scan

To further study the signaling pathways involved in Ca^2+^-mediated responses to salt in germinating seedlings, we analyzed 1,000 bp of the genomic sequence of presumed promoters of unique DEGs located upstream of each upregulated and downregulated gene that was responsive to NaCl, CaCl_2_, or the combined treatment for the presence of *cis*-elements, by an unbiased motif scan, using the Amadeus-Allegro tool (http://acgt.cs.tau.ac.il/allegro/download). The analysis of the presumed promoter sequences was performed to detect overrepresentation of *cis*-elements for TF binding. The JASPAR *cis*-element database (https://jaspar.genereg.net/) was used for motif detection. In response to the combined NaCl–CaCl_2_ treatment (Fig. 8), the unbiased motif analysis of upregulated responsive genes revealed a motif with consensus CACGTGTC (*P* value of 1.0E^-15^), which represents a G-box and an ABRE motif for the binding of bHLH TF family members, including TCPs, PIFs, SPT, MYC, and more. The second motif contained the consensus TCCGCGCA (*P* value of 6.0E^-15^), which is an alternative ABRE and CAMTA-binding motif (Kaplan et al., 2006; Finkler et al., 2007b). Analysis of these promoters using Pscan (http://159.149.160.88/pscan/) identified those TFs, including CAMTA1, CAMTA2 and CAMTA3 (full list of identified TFs under all conditions in Supplemental Table S5). Analysis of presumed promoters of genes downregulated in response to combined NaCl– CaCl_2_ treatment revealed two motifs with consensus ATGCTAAC (*P* value of 1.8E^-13^) and CCAGATAG (*P* value of 5.8E^-13^) (Fig. 8) that could not be verified as specific TF-binding elements. However, Pscan analysis identified *cis*-elements in these promoters that are considered to interact with multiple members of the TCP TF family (Supplemental Table S6). Presumed promoters of upregulated genes in response to NaCl treatment were found to harbor the consensus ACGTGTTCG (*P* value of 4.0E^-20^), which represents an ABRE and potential CAMTA-binding site (Kaplan et al., 2006; Finkler et al., 2007b; Shkolnik et al., 2019) and CTGCCGCG (*P* value of 2.7E^-18^), which represents a CGCG-core motif that is present in CAMTA-binding elements (Yang and Poovaiah, 2002; Han et al., 2006). In response to CaCl_2_ treatment, the presumed upregulated gene promoters were found to contain the consensus TAGGCCCA (*P* value of 5.1E^-23^), which represents a regulatory element group *cis*-element (Yamamoto et al., 2007) and GGTCCCGA (*P* value of 2.7E^-16^), which is classified as a *CYCLOIDEA(cyc)*-binding motif, has been shown to be involved in the regulation of floral development (He et al., 2022), and could be bound by TCP24, a TF that negatively regulates secondary cell-wall thickening (Wang et al., 2015). Downregulated presumed gene promoters identified in response to CaCl_2_ treatment were found to harbor the consensus GCCTGCAC (*P* value of 3.8E^-15^), reported to be the signature binding sequence of the nitrogen-control NtcA TFs in the unicellular cyanobacterium *Synechococcus* (Gunawardana and Herath, 2022), and a second motif with a consensus CGACGCGG (*P* value of 1.4E^-14^) which represents an ABRE and CAMTA-binding motif (Kaplan et al., 2006; Finkler et al., 2007b). The frequent representation of CAMTA-binding motifs in gene promoters in response to salt and Ca^2+^ coincides with the reported significant involvement of CAMTA TFs in the regulation of thousands of genes in response to environmental stimuli, including salt (Walley et al., 2007; Kim et al., 2013; Benn et al., 2014; Shkolnik et al., 2019).

**Figure 8.**
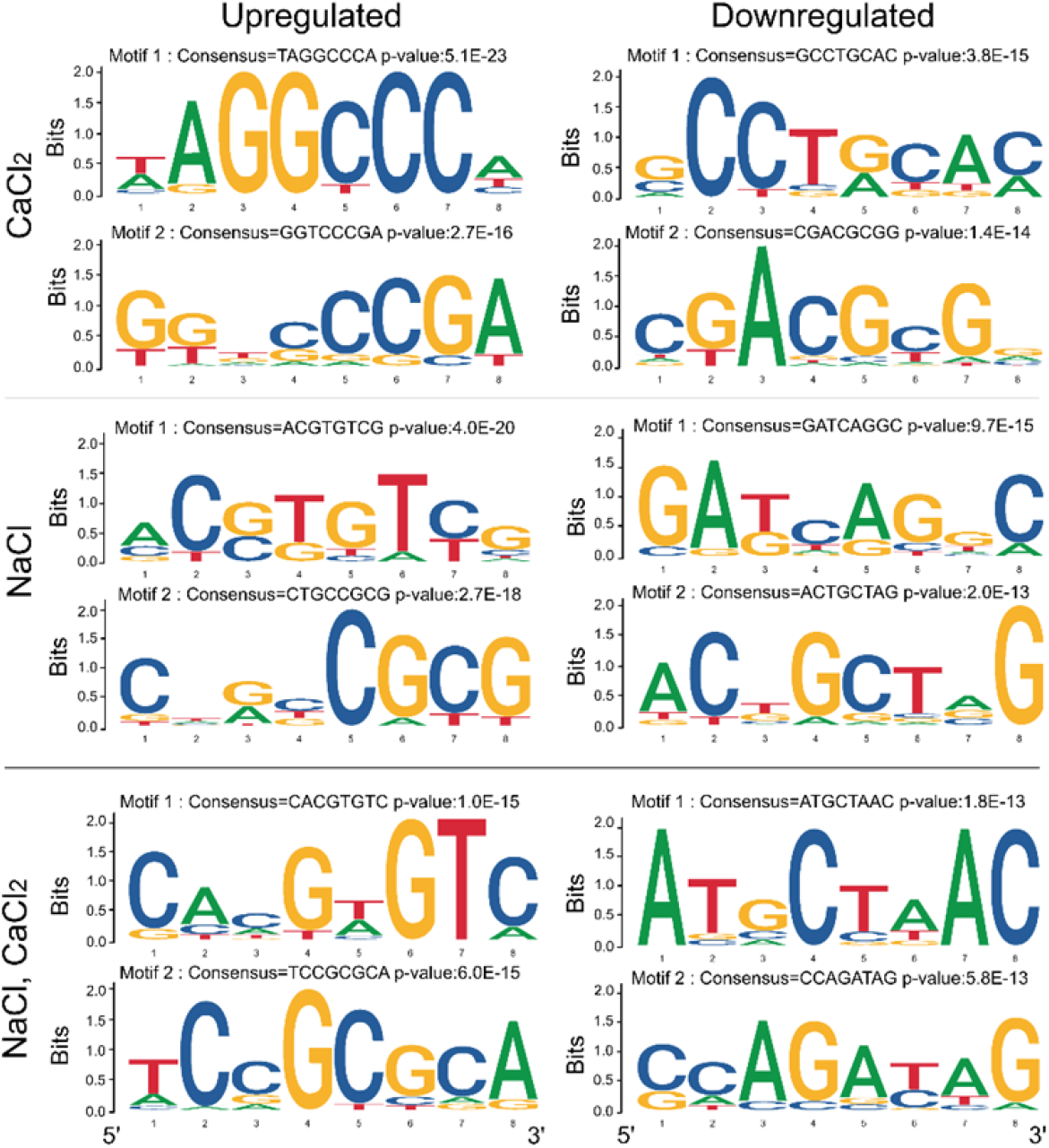
Highly represented sequence motifs in the promoters of differentially expressed genes which were identified as exclusively expressed in response to CaCl_2_, NaCl or combined CaCl_2_–NaCl treatments based on the data presented in Supplemental Table S4. Promoter analyses of differentially expressed genes were performed using Amadeus-Allegro software (http://acgt.cs.tau.ac.il/allegro/download.html) on the 1,000-bp genomic sequences upstream of the transcription start sites of the corresponding genes. Motifs of eight bases were derived from the JASPAR regulatory motif database. Left, promoter motifs in upregulated differentially expressed genes. In the promoters of unique genes upregulated by CaCl_2_ treatment, motif 1 with consensus TAGGCCCA and *P* value of 5.1E^-23^ represents a regulatory element group *cis*-regulatory element (Yamamoto et al., 2007). Motif 2 with consensus GGTCCCGA and a *P* value of 2.7E^-16^ is TCP24, a *CYCLOIDEA* (*CYC*)-binding motif known for regulation of leaf differentiation, floral development and regulation of secondary-wall thickening (Wang et al., 2015; He et al., 2022). In the promoters of unique genes upregulated by NaCl treatment, motif 1 with consensus ACGTGTTCG and a *P* value of 4.0E^-20^ represents an ABRE and CAMTA-binding site (Kaplan et al., 2006; Finkler et al., 2007b). Motif 2 with consensus CTGCCGCG and a *P* value of 2.7E^-18^ represents a CGCG-core motif that occurs in CAMTA-binding elements (Yang and Poovaiah, 2002; Han et al., 2006). In the promoters of unique genes upregulated by NaCl–CaCl_2_ treatment, motif 1 with consensus CACGTGTC and a *P* value of 1.0E-^15^ represents ABRE and CAMTA-binding site, and motif 2 with consensus TCCGCGCA and a *P* value of 6.0E^-15^ represents a CAMTA-binding motif (Kaplan et al., 2006; Finkler et al., 2007b). Right, promoter motifs in downregulated differentially expressed genes. In the promoters of unique genes downregulated by CaCl_2_ treatment, motif 1 with consensus GCCTGCAC and a *P* value of 3.8E^-15^ represents the signature binding sequence of NtcA transcription factors in cyanobacterium (Gunawardana and Herath, 2022), and motif 2 with consensus CGACGCGG and a *P* value of 1.4E^-14^ represents an ABRE and CAMTA-binding motif (Kaplan et al., 2006; Finkler et al., 2007b); the latter’s occurrence is not considered statistically significant. In promoters of unique genes downregulated by NaCl treatment, motif 1 with consensus GATCAGGC and a *P* value of 9.7E^-15^ has yet to be identified and motif 2 with consensus ACTGCTAG and a *P* value of 2.0E^-13^ has yet to be verified. In the promoters of unique genes downregulated by NaCl–CaCl_2_ treatment, two motifs were identified: ATGCTAAC (*P* = 1.8E^-13^) and CCAGATAG (*P* = 5.8E^-13^), which have yet to be verified.

## DISCUSSION

### HKT1;1 is involved in the Ca^2+^-mediated response to NaCl during germination

Seed germination is a critical developmental stage in the plant’s life cycle, when adjustments to environmental conditions, including salt stress, are necessary to ensure completion of germination and seedling establishment, as well as adequate vegetative development and yield. Although the positive effects of Ca^2+^ in improving germination and plant growth and development under salt-stress conditions have been demonstrated (Kent and Läuchli, 1985; Zehra et al., 2012; Shkolnik et al., 2019), our knowledge of the underlying mechanisms is fragmented. HKT1;1 is a Na^+^/K^+^ symporter that functions as an important factor in plant responses to salt stress (Rus et al., 2001; Laurie et al., 2002; Shkolnik-Inbar et al., 2013). Recently, *HKT1;1* was found to be regulated by the salt-regulated TF CAMTA6 during germination, contributing to maintaining Na^+^ homeostasis under salt stress (Shkolnik et al., 2019). These findings strongly suggest that HKT1;1 is a component of the Ca^2+^-signaling pathway in response to salt stress. Moreover, in the background of the *camta6* mutation, *HKT1;1* was expressed exclusively in the radicle of germinating seedlings, suggesting an association of this expression pattern with tolerance to salt during germination (Shkolnik et al., 2019). In *Arabidopsis*, maize, tomato and wheat, *HKT1* homologs have been found to promote tolerance to salt by facilitating Na^+^ retrieval from the root xylem sap to vascular parenchyma cells, thereby preventing Na^+^ accumulation in the shoot (Laurie et al., 2002; Sunarpi et al., 2005; Møller et al., 2009; Shkolnik-Inbar et al., 2013; Jaime-Pérez et al., 2017; Zhang et al., 2018). In contrast, the role of HKT1;1 during germination in the presence of NaCl, when the vascular tissues have not yet differentiated, has not been extensively investigated. Mutagenesis of *AtHKT1;1* and downregulation of *MtHKT1;1* and *MtHKT1;2* resulted in hypersensitivity to salt during germination, associated with increased accumulation of Na^+^ (Shkolnik et al., 2019; Zhang et al., 2019). We compared the germination rates of wild-type and two allelic *hkt1-*mutant seeds in the presence of NaCl and elevated CaCl_2_ concentration, and revealed lack of responsiveness of the *hkt1* mutants to CaCl_2_ supplementation compared to the wild-type, which displayed Ca^2+^-mediated improvement of germination rate under salinity (Fig. 1). Interestingly, CaCl_2_ application was found to promote radicle-focused expression of *HKT1;1*, which was further enhanced in the presence of NaCl (Fig. 2), and resembled the expression on the *camta6* background (Shkolnik et al., 2019). Hence, this radicle-focused expression of *HKT1;1* appears to correlate with improved germination rate in the presence of germination-arresting concentrations of NaCl. Collectively, the radicle-focused *HKT1;1* expression in the *camta6* background, which is not affected by NaCl application, the *camta6* salt-tolerance phenotype, the *camta6/hkt1* double mutant salt-hypersensitivity phenotype (Shkolnik et al., 2019), and hypersensitivity of *hkt1* which is not improved by CaCl_2_ supplementation (Fig. 1), all suggest the involvement of CAMTA6 in Ca^2+^-mediated regulation of *HKT1;1* expression under salt stress during germination. However, we still needed to address whether CAMTA6 directly regulates the transcription of *HKT1;1* by binding to its promoter. We assessed the possibility that the reduced expression of *HKT1;*1 in cotyledons in response to CaCl_2_ application, and not the radicle-focused expression, underlies the improved salt tolerance, by generating transgenic plants in which *HKT1;1* is expressed under the regulation of the promoter of the gene AT4G31830, whose expression is confined to the radicle (Jeong et al., 2014). Strikingly, this artificial radicle-confined expression of *HKT1;1* allowed germination in the presence of NaCl (Fig. 3B), similar to the effect of CaCl_2_ application. Therefore, the hypersensitivity to NaCl of *hkt1* mutants (Fig. 1; Shkolnik et al., 2019), in which functional HKT1;1 is absent from all of the germinating seedling’s tissues, and the improved tolerance gained by radicle-focused expression, indicate that this HKT1;1 expression pattern underlies, at least in part, the molecular mechanism of the Ca^2+^-mediated improvement in salt tolerance during germination. HKT1;1 is negatively regulated by the phosphatase PP2C49 (Chu et al., 2021). In accordance with that report and the *HKT1;1*-expression pattern presented herein (Fig. 2), *PP2C49* was predominantly expressed in the radicle of the germinating seedling and its expression was attenuated in response to CaCl_2_ treatment (Fig. 6), further indicating the association of *HKT1;1* expression in the radicle with the Ca^2+^-mediated response to salt during germination. Seeds of three allelic mutants of *PP2C49* displayed improved salt tolerance during germination compared to wild-type seeds (Fig. 5), in correlation with Ca^2+^-induced downregulation of *PP2C49* expression (Fig. 6, Supplemental Tables S1 and S4). Therefore, Ca^2+^ is suggested to downregulate the expression of *PP2C49*, resulting in attenuated inhibition of HKT1;1, which in turn functions to promote germination in the presence of NaCl by maintaining a low Na^+^/K^+^ ratio (Fig. 4).

### HKT1;1 is required for Ca^2+^-mediated maintenance of ion homeostasis under salt stress

Maintaining ion homeostasis at the cellular, tissue and whole-plant levels under salt stress is crucial for appropriate metabolism, physiology, growth and development of plants (Amin et al., 2021). A major harmful feature of salt stress in plants, caused by excessive Na^+^ levels, is ion toxicity, whereby the accumulation of noxious concentrations of Na^+^ in the cytosol impairs the essential conservation of cellular K^+^ level homeostasis (Zhu, 2003; Wang and Wu, 2013; Sun et al., 2015). With the exception of several C4 plants, Na^+^ is a non-essential element (Kronzucker et al., 2013; Nieves-Cordones et al., 2016). In contrast, K^+^ transport is critical for adequate enzyme activity, regulation of membrane potential, control of cytoplasmic and endosome luminal pH, and osmoticum level, which is required for cell-volume regulation (Zhu, 2003; Wang and Wu, 2013). Therefore, maintaining K^+^ and Na^+^ homeostasis is critical for salt-stress tolerance. Here, we sought to assess the role of HKT1;1 in maintaining Na^+^ and K^+^ homeostasis at early stages of seed germination, and the possible involvement of Ca^2+^ signaling in adjusting Na^+^/K^+^ ratios under salt stress. The previously demonstrated regulation of *HKT1;1* by the NaCl-regulated CAMTA6 (Shkolnik et al., 2019), whether direct or indirect, associates HKT1;1 with salt-responsive Ca^2+^ signaling. Moreover, pre-germinating *camta6* mutants were found to accumulate less Na^+^ in response to NaCl treatment, which was attributed to HKT1;1 and not to other key Na^+^ transporters, including SOS1 or NHX1 (Shkolnik et al., 2019). Ion-content analysis, performed to quantify Na^+^ and K^+^ concentrations in wild-type and *hkt1* pre-germinating seedlings, revealed a higher level of Na^+^ and a lower level of K^+^ in *hkt1* compared to wild-type seedlings treated with NaCl or NaCl and CaCl_2_ (Fig. 4). Interestingly, supplementation of CaCl_2_ along with NaCl significantly reduced the accumulation of Na^+^ while elevating K^+^ accumulation in wild-type but not *hkt1* seedlings, the latter being significantly K^+^-deprived (Fig. 4). Together, these findings indicate the requirement of a functioning HKT1;1 to facilitate the low Ca^2+^-regulated Na^+^/K^+^ ratios, reflected by the *hkt1* seedling’s hypersensitivity to NaCl during germination which is not rescued by CaCl_2_ application (Fig. 1). Finally, *pp2c49* mutants were found to maintain low Na^+^/K^+^ ratios under salt stress with or without CaCl_2_ supplementation (Fig. 5B), further indicating the involvement of PP2C49 in the Ca^2+^-mediated response to NaCl by negative regulation of HKT1;1 activity.

### Ca^2+^ modifies the transcriptome of germinating seedlings under salt stress

Exposure to salt stress significantly alters gene transcript-accumulation patterns in germinating and young *Arabidopsis* seedlings (Shkolnik et al., 2019; Sako et al., 2021). The broad influence of the *camta6* mutation on the transcriptomic profile of a germinating *Arabidopsis* seedling under control and salt stress conditions strongly indicates major involvement of Ca^2+^-signaling in the regulation of gene expression in response to salt (Shkolnik et al., 2019). Pscan analysis of over-represented TF binding-site motifs in the sequences of genes regulated by CaCl_2_, NaCl, or both combined provided TF lists to further understand the nature of Ca^2+^’s effect on the possible involvement of the identified TFs in the germinating seedling salt response. Presumed promoters of DEGs that were downregulated in response to the combined treatment contained *cis*-elements known to be bound by multiple TF members of the TCP family, including TCP2, TCP3, TCP4, TCP14, TCP17, TCP24 and BRC1 (Supplemental Table S6). These bHLH-containing DNA-binding motif (TCP domain) TFs are known to regulate plant development and defense responses by activating the biosynthetic pathways of bioactive metabolites, including brassinosteroids, jasmonic acid and flavonoids (Li, 2015). TCP24 is involved in the regulation of secondary cell wall-thickening processes (Wang et al., 2015). Conversely, analysis of presumed promoters of DEGs that were upregulated in response to the combined treatment revealed other involved TCPs, including TCP7, TCP8, TCP16, TCP20, TCP21, TCP23 and PTF1 (Supplemental Table S6), further suggesting a possible key role for TCPs in modulating the Ca^2+^-mediated response to salt during germination. Interestingly, *cis*-elements associated with the function of the ABA-regulated TFs ABI4 and ABI5, known to mediate ABA-controlled seed germination and developmental processes, including response to salt (Finkelstein, 1994; Finkelstein and Lynch, 2000; Shkolnik and Bar-Zvi, 2008; Skubacz et al., 2016; Maymon et al., 2022), were frequently found in the promoters of the DEGs that were upregulated in response to the combined treatment (Supplemental Table S6). Both ABI4 and ABI5 have been demonstrated as transcription co-activators but also as repressors of gene expression (Arenas-Huertero et al., 2000; Söderman et al., 2000; Shkolnik-Inbar et al., 2013; Huang et al., 2017). Seeds of *abi4* and *abi5* mutants display improved germination rate under salt stress (Quesada et al., 2000; Carles et al., 2002). Collectively, these data suggest that the Ca^2+^-signaling pathway involves regulation of the expression of ABI4- and ABI5-regulated genes to promote germination in the presence of salt. These data correspond with the functional classification analysis that presented enrichment of ∼12% of the DEGs that were downregulated in response to NaCl treatment as related to “response to ABA,” while ∼14% of the DEGs that were upregulated in response to CaCl_2_ treatment were classified in this category (Fig. 7, Supplemental Table S5). This classification feature, which was not characterized for DEGs in the combined treatment, possibly due to the two chemicals’ effects canceling each other out, depicts an opposing tendency of NaCl and CaCl_2_ on ABA-signaling pathways that suggests that Ca^2+^, along with the control of ion homeostasis, also attenuates the germination-arresting effect of ABA under salt stress. In addition, *cis*-elements that are associated with the bZIP TF HY5 and its homolog HYH, known for their involvement in light-mediated seedling development (Zhang et al., 2017), were identified in the presumed promoters of DEGs that were downregulated in response to CaCl_2_ treatment and in those of DEGs that were upregulated in response to NaCl or the combined treatment (Supplemental Table S6). This suggests that Ca^2+^ functions to adjust the expression of HY5- and HYH-regulated DEGs to allow germination in the presence of salt. Furthermore, *cis*-elements that are associated with the activity of members of the CAMTA protein family—CAMTA1, CAMTA2 and CAMTA3—were identified in the presumed promoters of DEGs that were downregulated by CaCl_2_ treatment and upregulated by NaCl or the combined treatment (Supplemental Table S6). These three CAMTAs have been reported to be involved in the regulation of thousands of genes in response to cold stress (Kim et al., 2013), and activation of CAMTA-repressed genes has been suggested to involve ubiquitin-mediated degradation of CAMTA, as in the case of CAMTA3 (Zhang et al., 2014). Support for the possible significant involvement of CAMTAs in response to CaCl_2_ and NaCl was provided by the motif scan performed using the Amadeus-Allegro tool, which revealed the ABRE and potentially CAMTA-binding *cis*-element motif CACGTGTC (Kaplan et al., 2006; Finkler et al., 2007b; Shkolnik et al., 2019) in the presumed promoters of NaCl- and CaCl_2_-upregulated DEGs (Fig. 8), in agreement with the reported involvement of CAMTA6 in regulating myriad genes in response to salt stress (Shkolnik et al., 2019). Notably, this specific full motif was identified as occurring four times in the *PP2C49* promoter, which was found to be downregulated by CaCl_2_ (Fig. 6, Supplemental Table S6), one in reverse-complement fashion, 287, 386 and 922 bp, and a single alternative ABRE and CAMTA-binding motif 408 bp upstream of the gene translation start codon (Supplemental Fig. S4). Therefore, CAMTAs are highly likely to function in modulating the complex response to CaCl_2_ in the presence of salt in pathways that remain to be deciphered.

## Acknowledgments

This research was supported by The Israel Science Foundation (grant No. 756/20 to D.S.).

## MATERIALS AND METHODS

### Plant materials, growth and stress assays

Arabidopsis (*Arabidopsis thaliana*) Col-0 plants were used in this research. The following mutants were obtained from the Arabidopsis Stock Center in Columbus, OH: *hkt1* (CS68521), *hkt1-4* (CS68092), *pp2c49-1* (SALK_111549C), *pp2c49-2* (SALK_015078C), and *pp2c49-3* (SALK_135177C). *HKT1;1pro*:GUS-expressing plants were generated by subcloning the 500- or 2,000-bp genomic DNA fragment upstream of the *HKT1;1* (At4G10310) gene translation start codon into pCAMBIA 1391Z binary vector at the *Sal*I and *EcoR*I sites upstream of the GUS-coding sequence, as previously described (Shkolnik et al., 2019). *PP2C49pro*:GUS-expressing transgenic plants were generated by subcloning the 1151-bp genomic DNA fragment upstream of the *PP2C49* (AT3G62260) gene translation start codon into pCAMBIA 1391Z at the *Sal*I and *EcoR*I sites. *AT4G31830pro*-HKT1;1_CDS_ (HKT1;1-RE)-expressing transgenic plants were generated for radicle-specific expression of HKT1;1, as previously described for the AT4G31830 expression pattern (Jeong et al., 2014). The construct was generated by introducing 692 bp of the promoter region of AT4G31830 into the pART27 binary vector recognition sites of *Sac*I and *Not*I. The *HKT1;1* coding sequence construct was generated by PCR with the total cDNA preparation as template which was synthesized using the High-Capacity cDNA Reverse Transcription Kit (Thermo Fisher Scientific) with oligo-dT as the primer, followed by PCR using Q5^®^ high-fidelity DNA polymerase (NEB) with specific primers (Table S2), and introduced into the vector’s *Not*I and *Spe*I sites; the Octopine Synthase (OCS) terminator was PCR-amplified using specific primers (Supplemental Table S7) and introduced into the *Spe*I and *Nhe*I sites. The construct was transformed into *hkt1-4* plants using the floral dip method (Clough and Bent, 1998) and transformants were selected based on resistance to kanamycin.

Seed surface sterilization and germination assays in the presence of different NaCl concentrations were performed as previously described (Shkolnik and Bar-Zvi, 2008). Germination on NaCl, CaCl_2_ or their combination, and CaCl_2_ or mannitol assays were performed in the same manner. A seedling was considered germinated when green fully unfurled cotyledons were observed. Treatments with NaCl prior to GUS staining, and RNA isolation for RNA-sequencing analysis and qRT-PCR were performed by seed imbibition at 4°C in the dark for 2–3 days prior to plating on Whatman filter paper (grade 1) soaked with 0.25X MS medium (Murashige and Skoog, 1962) and supplemented with the indicated chemicals. Pre-germinating seedlings were gently ejected manually, using curved fine forceps, 16 h after starting the treatment. GUS staining was performed as previously described (Weigel and Glazebrook, 2002), and images were taken using the Nikon SMZ18 stereoscope system equipped with a DS-Fi3 camera. Separation of cotyledon and radicle tissues for RNA isolation and subsequent qRT-PCR analysis was performed by relatively rough seedling ejection from the seed coat and tissue collection in separate test tubes using a fine pipette under a binocular. Treatment of several-day-old seedlings with NaCl, CaCl_2_, or both was performed by placing the seedlings such that the root is in contact with the agar-solidified 0.25X MS medium supplemented with the indicated chemicals, and the shoot is in the air to avoid direct contact with NaCl, as previously described (Shkolnik-Inbar and Bar-Zvi, 2012; Shkolnik et al., 2019).

### RT-qPCR analysis

Total RNA was isolated from pre-germinating seedlings using the ZR Plant RNA MiniPrep Kit (Zymo Research), and total cDNA was synthesized using the High-Capacity cDNA Reverse Transcription Kit (Thermo Fisher Scientific). The reaction mixture was prepared according to the manufacturer’s instructions, with random primers, and supplemented with 1 µg of total RNA. PCR mixtures (10 µL), containing 5 µL Fast SYBR Green Master Mix (Applied Biosystems by Thermo Fisher Scientific), 500 nM reverse and forward primers designed to amplify 80–120 bp of the genes of interest, and 20 ng cDNA were subjected to the Rotor-Gene Q 5-Plex HRM real-time PCR system (Qiagen), using the default program. The *PP2A* (AT1G69960) gene was used as an endogenous control. Relative quantification data were analyzed by the Rotor-Gene Q Pure Detection software V2.3.5 (Qiagen).

### Ion-content analysis

Control, NaCl-, CaCl_2_-, or NaCl–CaCl_2_-treated pre-germinating seedlings were ejected from seeds (see Plant materials, growth and stress assays) and samples for determination of Na^+^ and K^+^ content were prepared as previously described (Kalifa et al., 2004; Shkolnik-Inbar et al., 2013; Shkolnik et al., 2019). In each experimental repetition, 40 seedlings of each line were sampled. Ion content was determined using Arcos ICP-OES (Spectro/Ametek).

### RNA sequencing and bioinformatics analysis

RNA was isolated from ∼50 pre-germinated seedlings (three independent replicates, total n ∼150) of Col-0 that were treated without (control) or with 200 mM NaCl, 10 mM CaCl_2_ or both chemicals for 16 h (see Plant materials, growth and stress assays) using the ZR Plant RNA MiniPrep Kit (Zymo Research). RNA sequencing was performed at The Mantoux Institute for Bioinformatics of the Nancy and Stephen Grand Israel National Center for Personalized Medicine, Weizmann Institute of Science, Rehovot, Israel. Illumina RNA sequencing was performed in triplicate (Illumina Hiseq 2500; Illumina TruSeq RNA library Preparation Kit v2), resulting in 50-bp reads. Sequencing libraries were prepared using LIB_PREP_PROTOCOL. READ_TYPE reads were sequenced on NUM_LANES lane(s) of an Illumina ILLUMINA_INSTRUMENT. The output was ∼18 million reads per sample. Mapping and alignment were performed using TopHat version 2.1.1 and Bowtie2 version 2.5.1 (reference genome TAIR10), followed by gene counting performed by HTseq-count version 2.0.3. Values above 10 were used for normalization and gene-expression analysis (DESeq2 version 3.17). Partek Genomics Suite was used for further analysis (version 7.20.083). Functional classification of DEGs was performed with the DAVID Gene Functional Classification tool (https://david.ncifcrf.gov/). The tool was used with multiple database sources of functional annotations. Enriched significant classifications were selected based on *P* values (*P* ≤ 0.05). Enrichment of the upregulated and downregulated DEGs was classified by biological process, molecular function and cellular component. Venn diagrams were created using the Venny tool (http://bioinfogp.cnb.csic.es/tools/venny/index.html; Oliveros, 2015). The Amadeus-Allegro tool (http://acgt.cs.tau.ac.il/allegro/download.html; Linhart et al., 2008; Orenstein et al., 2012) was used for motif analysis. Motif enrichment was performed on 1,000bp of the genomic sequences upstream of the presumed transcription start sites of the DEGs. Eight-base-long motifs were searched for based on the JASPAR regulatory motif database (https://jaspar.genereg.net/matrix/MA0970.1/).

### Statistical analysis

Data were analyzed using JMP software version pro16 and Microsoft Excel 2019 ToolPak.

### Accession numbers

Accession numbers of the major genes investigated in this research are: *HKT1;1*, AT4G10310; *PP2C49*, AT3G62260.

## Supplemental material for

### Supplemental figures

**Figure S1.**
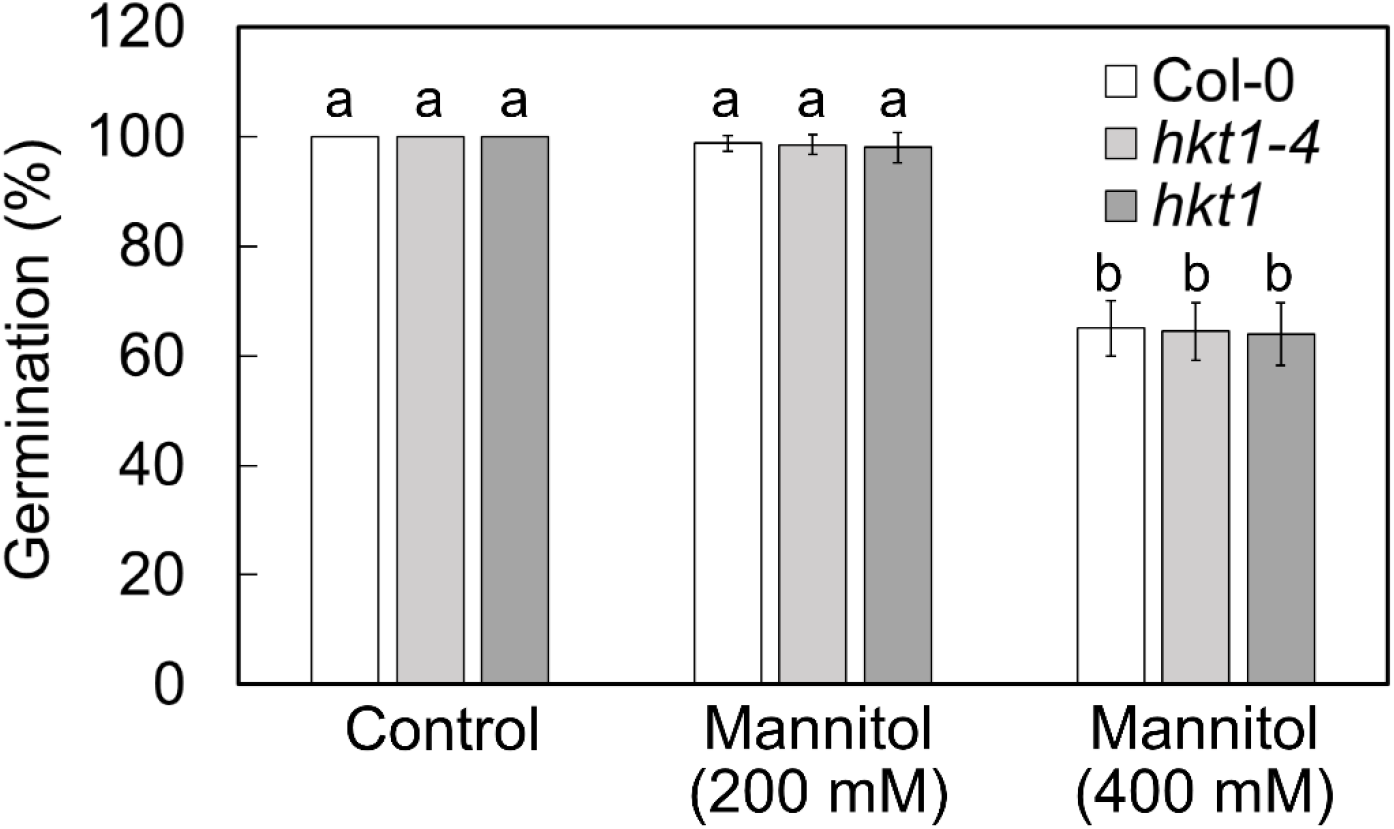
Germination assay on mannitol-containing media. Seeds of Col-0, *hkt1-4* and *hkt1* were plated on medium supplemented with the indicated concentrations of mannitol. Germination was scored 5 days later. Data are means ± SE (three independent biological experiments, ∼100 seeds each) and different letters above the bars indicate significantly different values by Tukey’s HSD post hoc test (*P* < 0.01).

**Figure S2.**
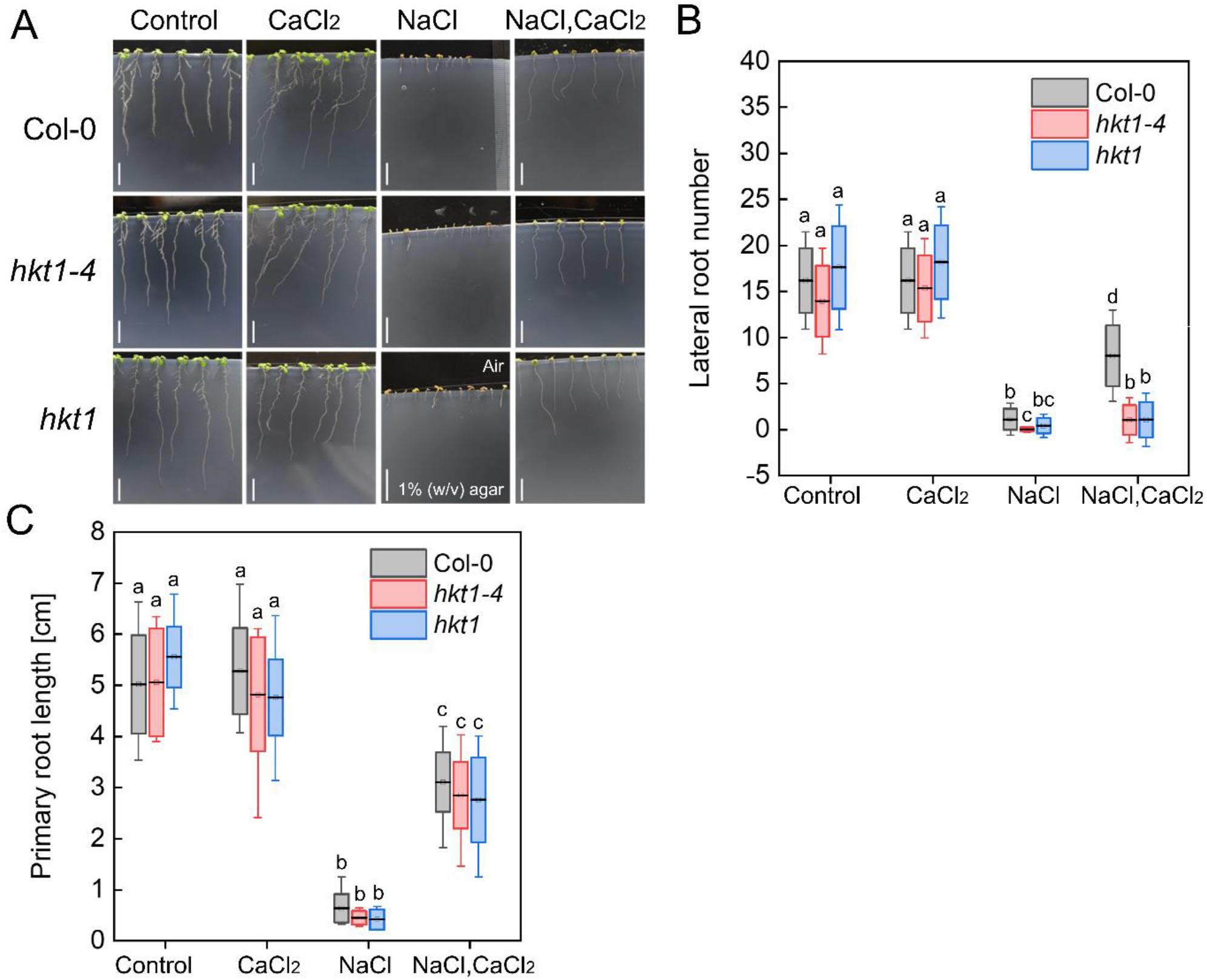
Root development phenotyping of *hkt1* mutants in response to NaCl, CaCl_2,_ or combined treatments. A, Three-day-old Col-0, *hkt1-4* and *hkt1* seedlings were transferred onto agar-solidified 0.25X MS medium (see Materials and Methods in the main text) supplemented with 0 (control), NaCl (200 mM), CaCl_2_ (10 mM), or both, as indicated. Bar = 1 cm. B, Lateral root number was scored using a stereoscope. C, Primary root length quantification. In B and C, data are mean values ± SD (three independent experiments, 8 seedlings each) and different letters above the bars indicate significantly different values by Tukey’s HSD post hoc test (*P* < 0.01).

**Figure S3.**
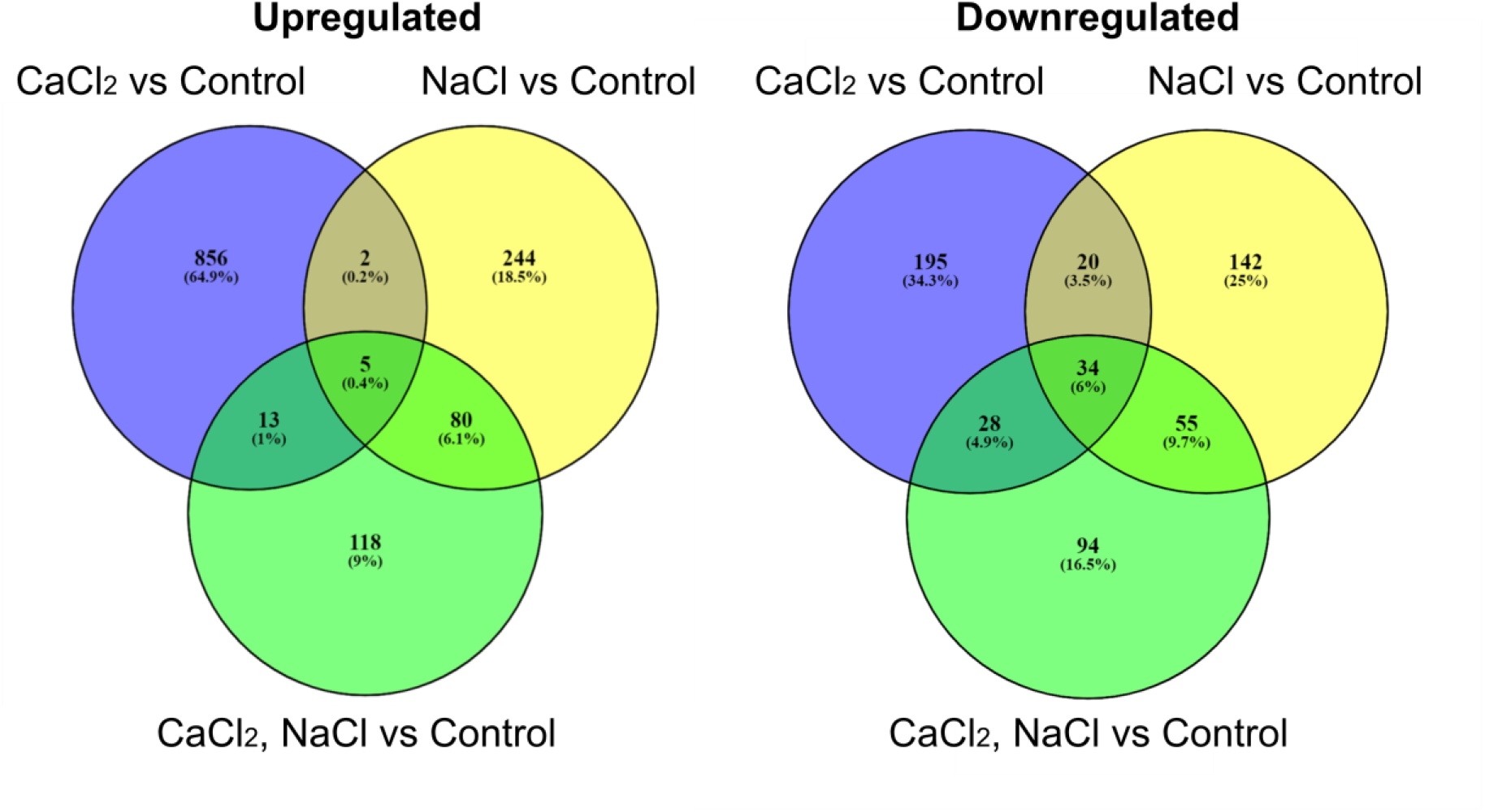
Venn diagrams of differentially salt-responsive genes. Venn diagrams were created to compare differentially expressed gene lists (cutoff FC ≥ |1.25| and *P* value < 0.05) using Venny tool: http://bioinfogp.cnb.csic.es/tools/venny/index.html (Oliveros, 2015), an interactive tool for comparing lists with Venn’s diagrams. The upregulated or downregulated gene groups identified by comparing gene lists of CaCl_2_, NaCl or CaCl_2_–NaCl treatments with the control treatment are indicated. CORRECT TO Upregulated, Downregulated

**Figure S4.**
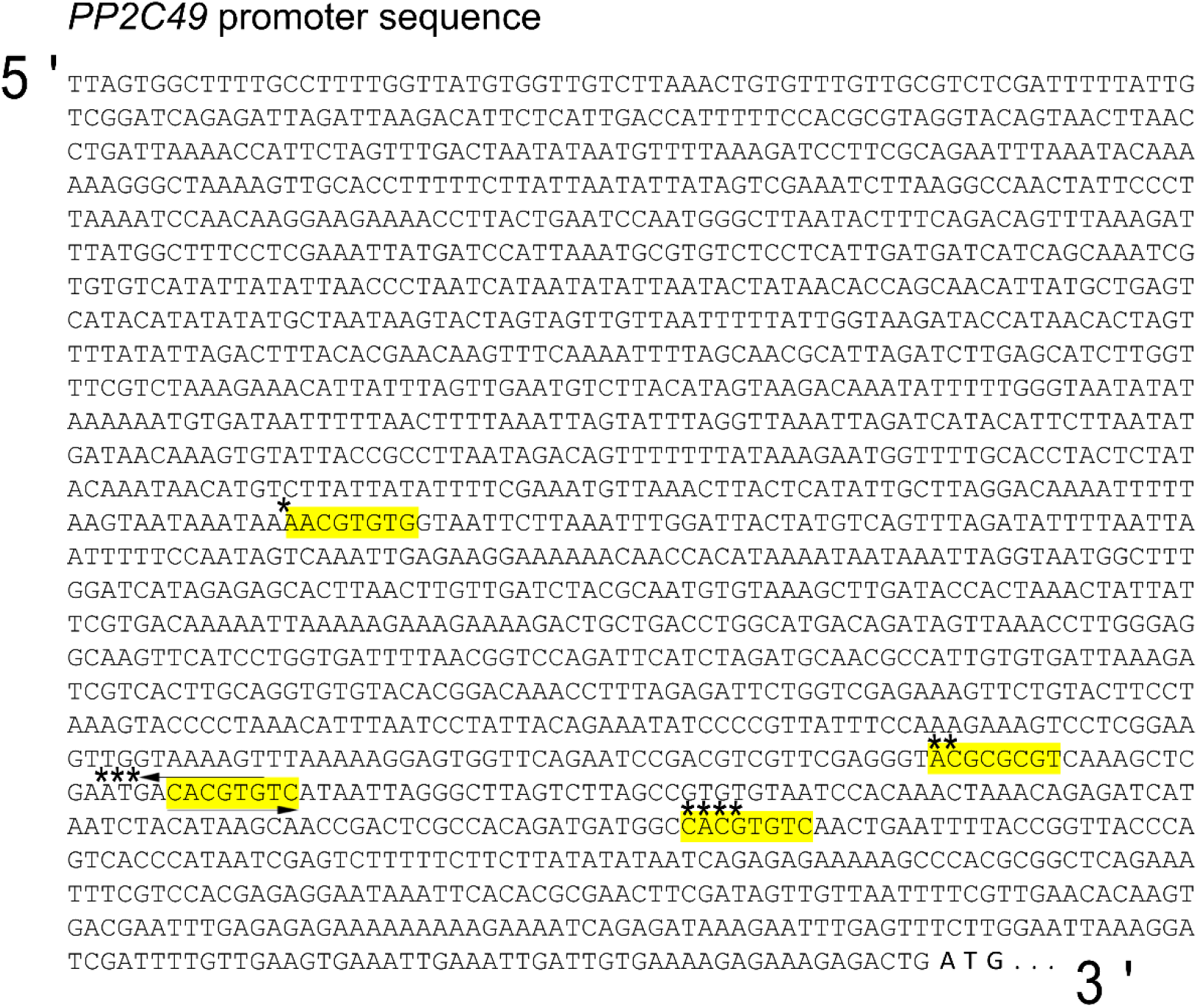
*PP2C49* promoter sequence as presented in TAIR10 database. Four ABRE and potential CAMTA-binding motifs were identified, *922 bp, ***386 bp (reverse-complement motifs are indicated with arrows), ****287 bp, and a single alternative ABRE and CAMTA-binding motif **408 bp upstream of the gene translation start codon (Kaplan et al., 2006; Finkler et al., 2007).

**Table S1.** List of differentially expressed upregulated and downregulated genes for CaCl_2_ treatment vs. control, following cutoff FC ≥ |1.25| and *P* value < 0.05, performed using Partek Genomics Suite hierarchical clustering.

**Table S2.** List of differentially expressed upregulated and downregulated genes for NaCl treatment vs. control, following cutoff FC ≥ |1.25| and *P* value < 0.05, performed using Partek Genomics Suite hierarchical clustering.

**Table S3.** List of differentially expressed upregulated and downregulated genes for CaCl_2_– NaCl treatment vs. control, following cutoff FC ≥ |1.25| and *P* value < 0.05, performed using Partek Genomics Suite hierarchical clustering.

**Table S4.** Lists of differentially expressed genes obtained by the Venny tool: http://bioinfogp.cnb.csic.es/tools/venny/index.html (Oliveros, 2015) with cutoff FC ≥ |1.25| and *P* value < 0.05 as presented in Figure S3 Venn diagrams.

**Table S5.** Functional classification of NaCl and/or CaCl_2_-responsive genes obtained by DAVID Gene Functional Classification tool (https://david.ncifcrf.gov/) (Sherman et al., 2022), with multiple sources of functional annotations (https://david.ncifcrf.gov/content.jsp?file=update.html). Classifications were selected based on *P* values (*P* ≤ 0.05). The enrichment analysis of the differentially expressed up- and downregulated genes was classified based on the following: biological process, molecular function and cellular component.

**Table S6.** Promoter scan analysis for transcription factor-binding sites in the upregulated and downregulated promoters of CaCl_2_ / NaCl / CaCl_2_–NaCl treatment vs. control using Ps

**Table S7.**
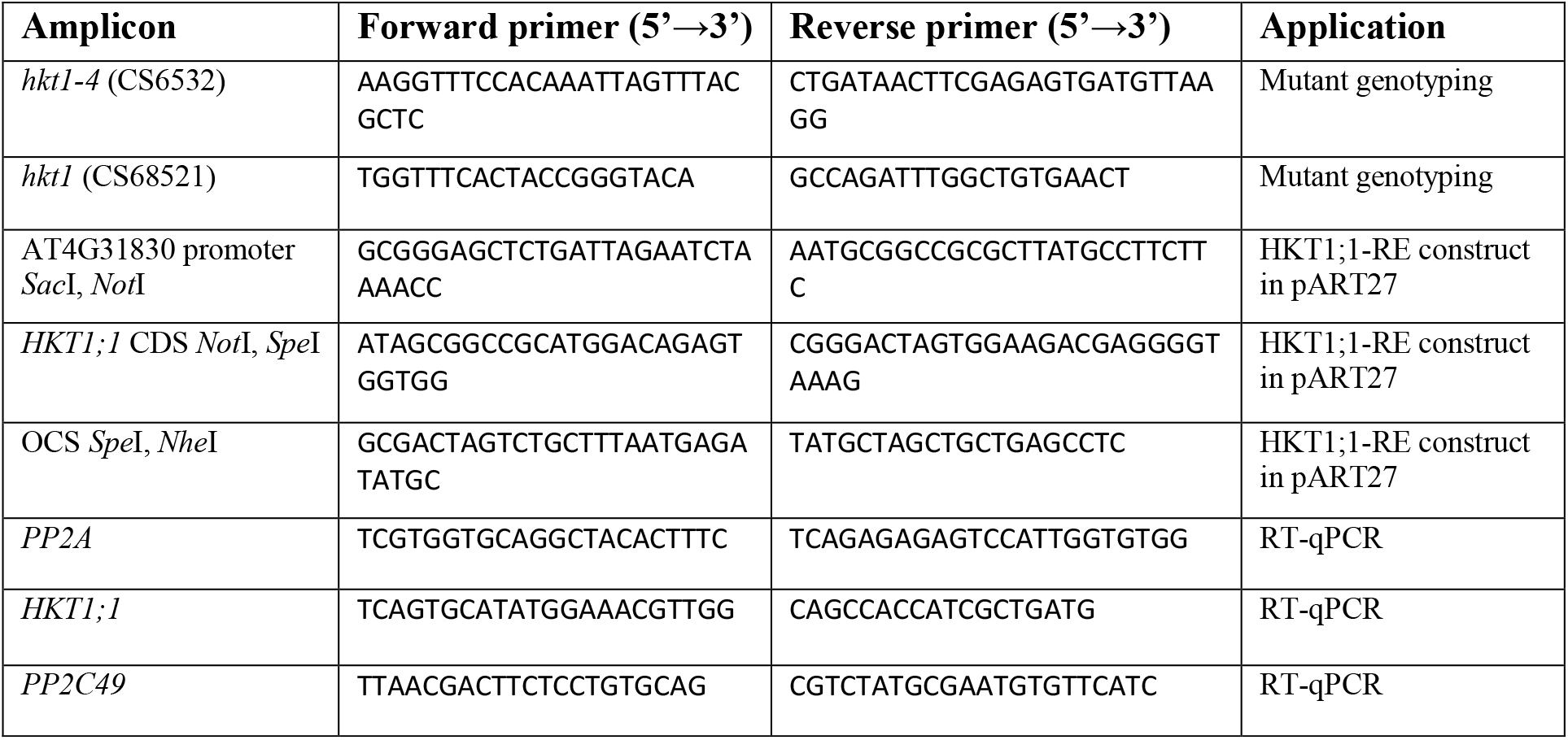
Primer list.

